# Synthetic Control of Implanted Engineered Liver Tissue Growth

**DOI:** 10.64898/2025.12.10.693527

**Authors:** Amy E. Stoddard, Vardhman Kumar, Constantine N. Tzouanas, Veronica Hui, Jeffrey Li, Anisha Jain, Alanna Farrell, Sangeeta N. Bhatia, Christopher S. Chen

## Abstract

Despite the promise of engineered tissue implants for the treatment of organ failure, scaling of these constructs to sizes of therapeutic relevance remains a barrier to clinical translation. Here, we propose a strategy to circumvent this limitation: to instead implant a small-scale construct and then induce it to grow *in situ* after its engraftment into a host. Using engineered liver tissue as a proof-of-concept application, we integrated synthetic biology and tissue engineering tools to build liver tissues that can be expanded on-demand after implantation *in vivo*. To achieve this goal, we first identified the combination of YAP and growth factor signaling as sufficient to drive human hepatocyte proliferation in dense, 3D engineered tissues. We then engineered control of these signaling axes using synthetic biology tools to drive human liver tissue expansion both *in vitro* and *in vivo*. As such, this work establishes a genetic strategy for generating large organ implants through bioengineered, on-demand outgrowth using synthetic triggers (BOOST).

**Teaser:** Presenting bioengineered on-demand outgrowth via synthetic biology triggering (BOOST) for *in situ* solid cell therapy scale up.

## Introduction

Organ transplant is currently the only curative treatment for patients with end stage organ failure, yet this therapy is inaccessible to many due to the paucity of organs available for transplant. Tissue engineers have made substantial advancements towards engineering tissue-based cell therapies which could serve as alternatives or bridges to transplantation (*1*, *2*). However, across a variety of applications, including engineered cardiac, kidney, muscle and liver tissue, the maximum size of the engineered tissue construct, and thus the maximum deliverable therapeutic dose, remains limited(*3*, *4*). In addition to the challenge of sourcing sufficient cellular raw materials for large constructs, assembly of these cells into a viable, large-scale implant remains unsolved(*5–7*). Despite innovations by the community in vascularization(*8–10*), bioreactor design(*11*, *12*), and complex assembly methods like 3D bioprinting(*13–16*), a reliable strategy for the assembly and perfusion of large or dense tissue implants remains elusive. To circumvent these limitations, we approached this fabrication challenge from a different angle, and instead asked if it would be possible to first implant a small-scale construct and then drive it to expand *in situ* after its engraftment into the host.

A key first step towards this method of *in situ* scale up would be the control of cellular growth within the engineered construct after engraftment. Previous studies primarily in the rodent heart and liver have explored the concept of triggering tissue growth and regeneration by delivering growth or transcription factors(*17–21*). These strategies, however, may be difficult to translate clinically because they rely not only on stimulation of proliferation, but also on resolution of the underlying structural damage of the diseased organ(*22*). Synthetic biologist have also applied such a framework to immunologic cell therapies like CAR T-cells, leveraging innovative genetic tools to precisely and inducibly control cell growth, activity, and death of the cell product(*23–26*). Inspired by these advances, we sought integrate these synthetic biology tools into tissue engineering to develop a strategy to induce <u>B</u>ioengineered <u>O</u>n-demand <u>O</u>utgrowth via <u>S</u>ynthetic biology <u>T</u>riggering (BOOST) of a tissue construct.

An ideal use case for this scale-up by growth strategy is the treatment of liver failure, as end stage liver disease is particularly fatal and can only be cured by organ transplant(*27*). As an alternative to transplant, injection of human hepatocytes (HEPs), the main functional cells of the liver, was explored, but ultimately has been clinically unsuccessful due to insufficient engraftment of the injected cells(*28*, *29*). Recent work by our group and others have demonstrated that instead, assembly of HEP aggregates into vascularized 3D tissues could enable their engraftment into ectopic sites, where they can provide long term functional support to the recipient(*30–32*). The scale up of these tissue therapeutics to human scale, however, remains a challenge, and would benefit from BOOST. To realize on-demand *in situ* scale up of engineered liver tissue, the right set of signals sufficient to promote human HEP proliferation, specifically in a 3D engineered human tissue context, would have to be identified. While these cues have yet to be fully defined, integration of insights from recent successes in *in vitro* HEP expansion(*33–37*), combined with decades of research on liver regeneration(*38*), provide clues as to the critical signaling axes that control HEP proliferative state.

In this study we leveraged 2D and 3D human liver culture models to identify, and then synthetically control, a set of signals sufficient to induce the proliferation of human HEPs in implanted 3D engineered hepatic tissues. We report that while GF and YAP signaling on their own are insufficient to stimulate the proliferation of HEPs in dense cultures, their concurrent activation synergizes to drive a robust proliferative response in human HEPs even in dense 3D, engineered tissues. Using synthetic biology tools to install local control over YAP and growth factor signaling, we demonstrate on-demand growth of engineered human liver constructs following their implantation into mice. As such, this work forms the foundation of an alternative strategy for achieving therapeutic-scale cell therapies: using synthetic biology to drive the expansion of engineered tissues after implantation.

## Results

### YAP and GF signaling synergize to induce proliferation of dense cultures of human hepatocytes

Towards our overall goal of controlling liver implant growth, we first set out to identify which signaling axes are sufficient to promote proliferation of HEPs in 3D engineered tissues. Primary human HEPs were seeded at a concentration of 3E4 or 3E5 cells/cm2 on collagen coated well plates, allowed to adhere overnight, and then treated with a panel of GFs and other compounds previously shown to promote HEP proliferation for 2 days (Fig. 1A). We found that in the absence of supplemental GFs, HEPs in a basal media (Advanced DMEM + B27 + P/S) did not proliferate. Of the 17 compounds tested, only HGF and the EGFR ligands, EGF and TGFα, were sufficient to robustly increase HEP proliferation (Fig. 1B, S1A). We selected HGF and TGFα for future study, as these compounds were the most strongly mitogenic in our platform. Interestingly, we did not observe a proliferative response to WNT ligands. Nevertheless, given the pivotal role for WNT in HEP expansion protocols and liver regeneration, we also selected WNT2 and RSPO3 (the most highly expressed WNT ligands in the human liver) for subsequent experiments. Having identified a set of GFs that were mitogenic to human HEPs, we were curious if these GFs could also stimulate proliferation of our engineered 3D liver tissues, where HEPs are arranged in dense co-aggregates with fibroblasts (FBs) and suspended in a fibrin hydrogel with endothelial cells (ECs) (Fig. 1C). Contrary to our prior findings, we found that HEPs in 3D liver tissues did not proliferate in response to a 1 week treatment with GFs (Fig. 1D). To determine if this observation was a result of density-mediated contact inhibition, which regulates most adherent cells, we compared the proliferative response to GFs in sparsely- versus densely-seeded HEP monolayers, and found that the GF-mediated proliferative response was lost in dense HEP cultures (Fig. 1E, S1B,C). Curious if this phenomenon was broadly applicable to a variety of mitogenic growth factor cocktails, we treated HEPs with media formulations from previously published 2D and 3D organoid expansion protocols(*33*, *36*) and found that a density dependent reduction in proliferation occurred in all cases (Fig. S1D).

**Figure 1.**
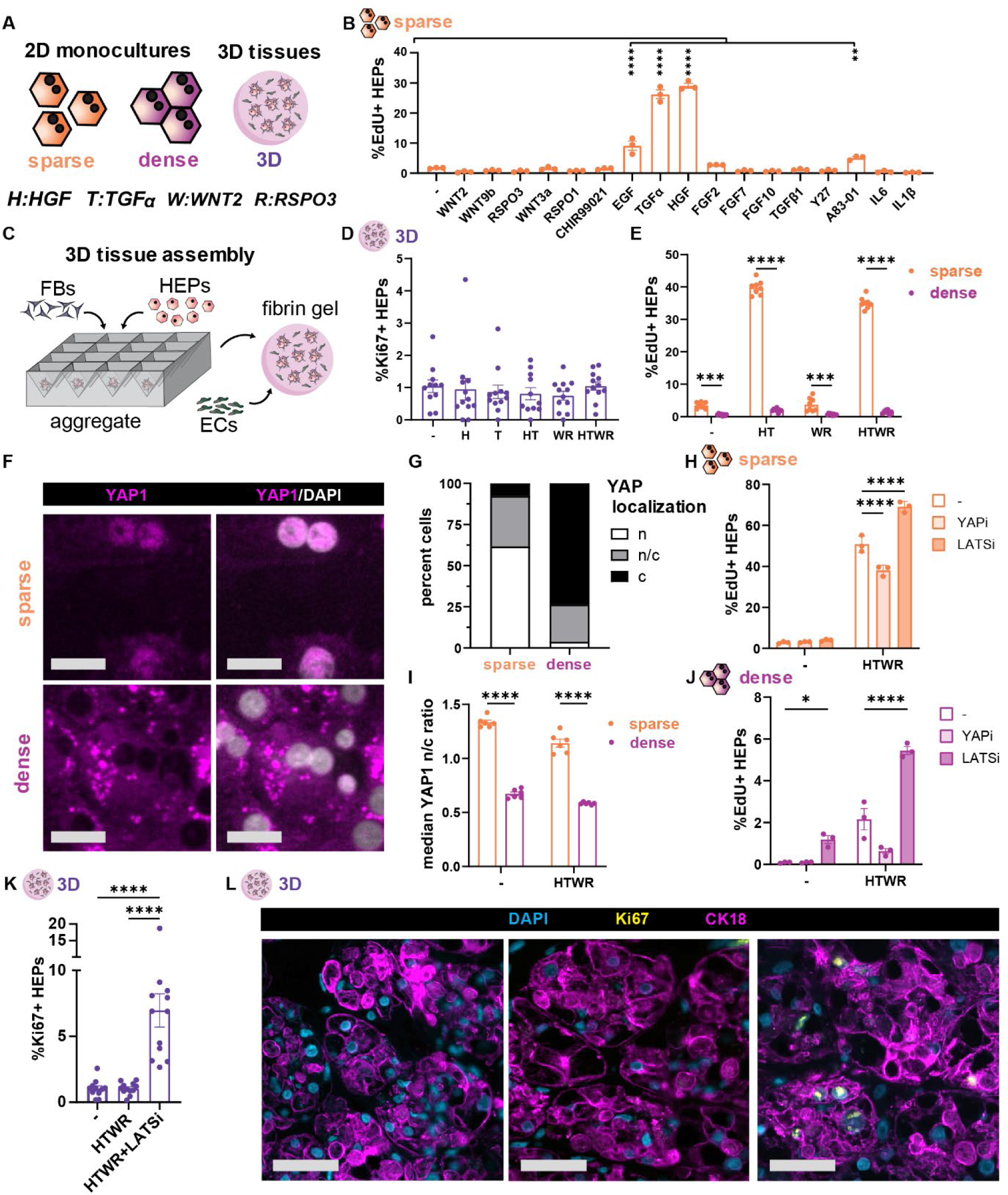
YAP and GF signaling synergize to induce proliferation of dense cultures of human hepatocytes. **(A)** Icon legend and factor abbreviations. **(B)** Proliferation of 2D sparse HEPs after 2 days of factor treatment in serum free media (N:3,n:3). **(C)** Schematic of 3D liver tissue assembly. **(D)** Proliferation of 3D liver tissues after 1 week treatment with combinations of recombinant GFs (N:3,n:4). **(E)** Proliferation of HEPs in response to recombinant GFs in sparse or dense 2D culture conditions (N:3,n:3).**(F)** Immunofluorescent staining for YAP in sparse and dense 2D cultures. Scale bars 25 µm **(G)** Binned YAP localization (n:nuclear/ c:cytoplasmic) of 2D dense and sparse HEPs in serum free media without supplemental GFs(N:2,n:3). **(H)** Median YAP nuclear to cytoplasmic ratio in 2D sparse and dense cultures, with and without GF supplementation (N:2,n:3). **(I)** Proliferation of sparse and **(J)** dense cultures with and without treatment with growth factors, and YAP inhibitor CA3, or LATS inhibitor TDI-011536 (N:3,n:3). **(K)** Proliferation of engineered 3D liver tissues treated with basal media, GFs, or GFs and LATS inhibitor (N:3,n:4). **(L)** Representative images from K, where HEPs are labeled with Arginase 1 (Arg1). Scale bars 50 µm. (p *<.05, ** < .01, *** < .001, **** < .0001; B, K: one-way ANOVA, E,I,H,J: two-way ANOVA; B,G,I,J: data shown from one representative experiment, see supplement for additional repeats).

To understand what proliferative regulators might be affected by cell density, thereby preventing GF-mediated stimulation of growth, we examined classical regulators. We found that YAP, a key transcriptional effector in the Hippo signaling pathway responsible for regulating cell proliferation and organ size(*39*), is excluded from the nucleus in dense cultures (Fig. 1F,G, S1E). Notably, YAP translocated to the nucleus in all sparse conditions tested, even in the absence of serum or growth factors, while YAP remained cytoplasmically-restricted in all dense cultures, even those treated with a mitogenic cocktail sufficient to promote proliferation of sparsely seeded HEPs (Fig. 1H, S1F,G,H). This density dependent reduction in YAP activity was further confirmed by a panel of YAP target genes (Fig. S1I), wherein 3D culture most strongly suppressed transcription of YAP target genes. As such, YAP signaling appears to be largely regulated by cell density and not by mitogenic GFs in human HEPs, suggesting that YAP imposes a density checkpoint distinct from GF regulation. To further explore this interplay, we independently modulated GF exposure and YAP activity and assessed the effects on HEP proliferation. In sparse cultures, where YAP is active, we find that partial inhibition of YAP by treatment with the YAP inhibitor CA3 resulted in a reduction of GF-mediated proliferation (Fig. 1I, S2A,C). Conversely, activation of YAP using LATS kinase inhibitor TDI-011536 (LATSi) in sparse cultures further increased proliferation, but was not sufficient to induce proliferation in the absence of GF co-stimulation (Fig. 1I, S2B,C). In dense cultures, where YAP is inactive, YAP activation via LATSi increased proliferation especially when combined with GF treatment (Fig. 1J, S2D). This effect extended into 3D; YAP activation also increased proliferation of HEPs in 3D engineered liver tissues (Fig. 1K,L). Together, these studies suggest that while neither YAP nor GF signaling is sufficient on its own to induce proliferation, the simultaneous activation of both pathways synergizes to robustly drive proliferation of human HEPs, even in dense 2D and 3D cultures that are typically growth arrested by contact inhibition.

### Synthetic control of GF and YAP signaling enables on demand proliferation of engineered 3D liver tissues

Having defined a set of cues sufficient to trigger HEP growth, we next sought to synthetically control these pathways locally within the implant, as systemic induction of proliferative signals is not a clinically feasible strategy for expansion of implanted tissues. We set out to engineer a synthetic biology toolkit capable of locally modulating growth factor and YAP signaling within engineered liver tissue, enabling on-demand control of proliferation even after implantation. To exert control over GF signaling, we engineered the FBs, which are present in our engineered liver tissues to support the HEPs through paracrine and cell contact-related interactions, to secrete HGF, TGFα, WNT2, or RSPO3 (or synthetic growth factors, synGFs) under doxycycline-inducible control (Fig. 2A). We validated the inducibility of cells and bioactivity of the protein products through a series of western blots, ELISAs, and conditioned media studies (Fig. 2B,C, S3A,B,C). We further confirmed that upon treatment with conditioned media from GF-producing FBs, HEPs proliferated in a manner consistent with our prior studies using recombinant GF (Fig. 2D, S3D). To control YAP activity, we directly engineered the HEPs in our tissues to express a degradation-deficient YAP mutant, YAP5SA (*40*), which signals constitutively (Fig. 2E). Using lentivirus, we found that HEPs could be transduced to express YAP5SA with an approximately 50% infection rate, and successful DOX dependent expression of the YAP5SA protein was confirmed (Fig. 2F). To further assess the activity of this gene product, we aggregated YAP5SA HEPs and found that DOX treatment increased the expression of the canonical YAP target gene CTGF by roughly 10-fold (Fig. 2G). Of note, this effect was most pronounced after 4 days of DOX induction as opposed to shorter treatment durations, suggesting a requirement for extended YAP5SA induction for robust effects on downstream gene expression.

**Figure 2.**
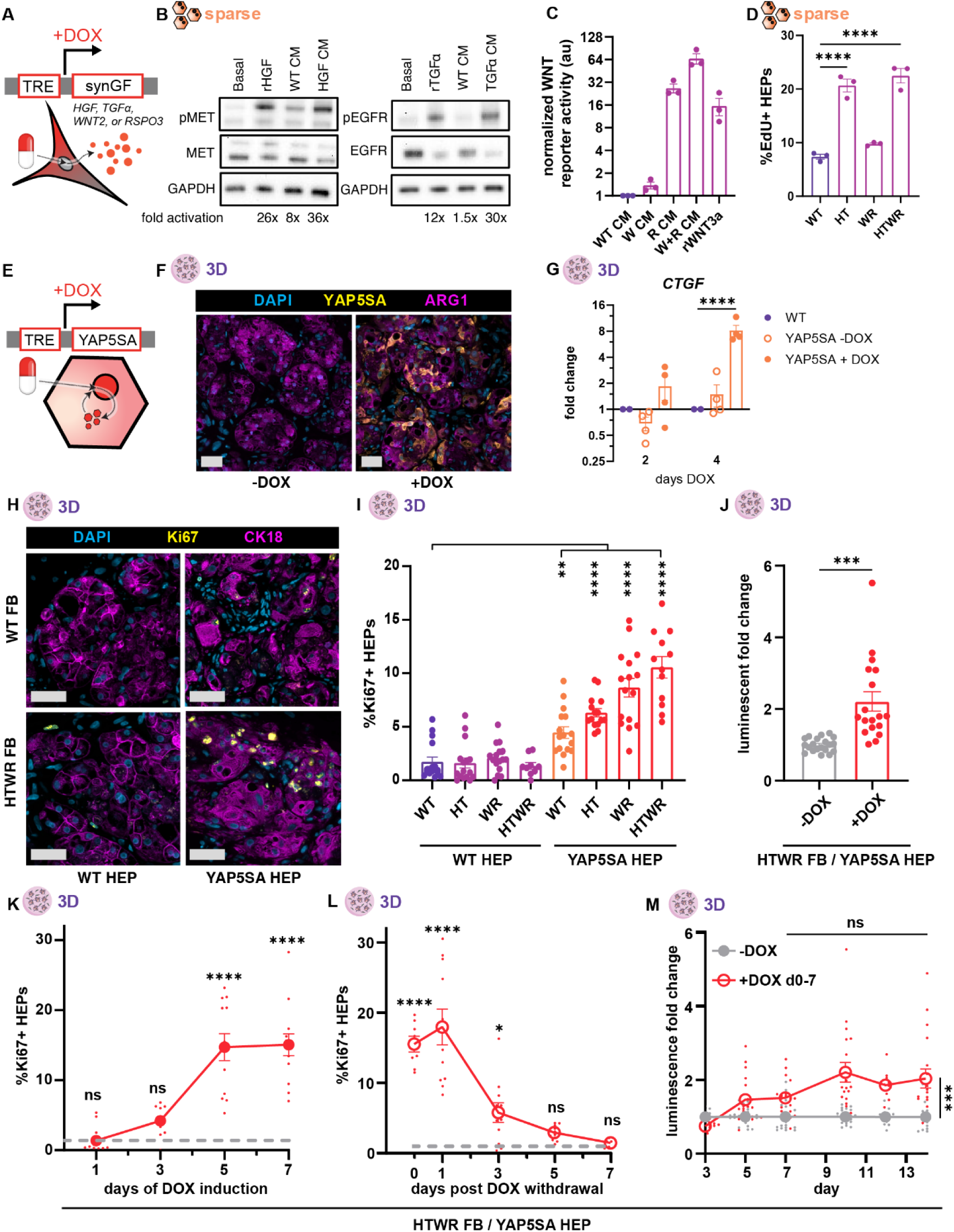
Synthetic control of GF and YAP signaling enables on demand proliferation of engineered 3D liver tissues. **(A)** FBs engineered to express GFs under doxycycline control. **(B)** Western blot for EGFR and MET activation in 2D HEPs treated with recombinant HGF or TGFα or conditioned media (CM) from HGF or TGFα producing FBs. **(C)** WNT reporter activity upon treatment with conditioned media (CM) from WNT2 or RSPO3 producing FBs or 100 ng/mL recombinant WNT3A (N:1,n:3). **(D)** Proliferation of 2D sparse HEPs treated with conditioned media from WT or synGF NHDFs for 2 days (N:2,n:3). **(E)** HEPs engineered to express YAP5SA under doxycycline control. **(F)** Immunohistochemical micrographs of HEPs in 3D engineered liver tissues stained for YAP5SA with and without DOX treatment. Scale bars 50 µm. **(G)** qPCR expression of YAP target gene CTGF by 3D WT or YAP5SA HEPs (+/- DOX) after 2 or 4 days of DOX induction (N:2,n:2). **(H)** Representative immunofluorescence staining of 3D tissues after 7 days of synGF and/or YAP5SA expression. Scale bars 50 µm **(I)** Quantification of HEP proliferation in 3D tissues after 7 days of synGF/YAP5SA expression (N:4,n≥3). **(J)** Luminescent reporter signal of FB^HTWR^ /HEP^YAP5SA^ tissues after 7 days of DOX induction compared to uninduced controls measured on day 10 (N:4,n≥4). Kinetic curve of proliferative response to DOX induction **(K)** and to DOX withdrawal after 7 days DOX induction **(L)** by HEPS in FB^HTWR^ /HEP^YAP5SA^ tissues. Uninduced proliferation baseline denoted by grey dotted line (N:3,n:4). **(M)** Luminescent reporter signal of FB^HTWR^ /HEP^YAP5SA^ tissues induced with DOX from day 0 to day 7 compared to uninduced controls (N:4,n≥4). (p *<.05, ** < .01, *** < .001, **** < .0001; J: Welch’s t-test, C,I,K,L: one-way ANOVA, G,M: two-way ANOVA; D: data shown from one representative experiment, see supplement for additional repeats).

After confirming GF and YAP5SA construct efficacy, we assembled engineered liver tissues from combinations of wild-type or GF-expressing FBs and wild-type or YAP5SA-expressing HEPs. Consistent with our findings using recombinant growth factors, we found that synGF production alone did not increase proliferation of unmodified HEPs in 3D liver tissues, regardless of GF combination (Fig. 2H,I). Similarly, YAP5SA expression in HEPs without concurrent synGF induction caused only marginal increases in HEP proliferation. Combining both synGF FBs and YAP5SA HEPs, however, triggered a profound synergistic enhancement of HEP proliferation. The strongest proliferative response occurred with the combination of all four GFs and YAP5SA (FB^HTWR^ /HEP^YAP5SA^), resulting in an approximately 6-fold increase in proliferation compared to unengineered (FB^WT^/HEP^WT^) tissues. Using an integrated luminescent reporter, we find that FB^HTWR^ /HEP^YAP5SA^ tissues have twice as many engineered HEPs as controls (Fig. 2J). This expansion was similarly reflected by a 50% increase in HEP nuclei and CK18+ HEP area, consistent with a doubling of the engineered HEPs in response to a 7-day DOX stimulus (Fig. S3E,F).

For clinical safety, it is critical that synthetically induced proliferation is triggered only by treatment with the small molecule activator, and that this induction ceases upon inducer withdrawal. To confirm the inducibility of our system, we compared FB^HTWR^ /HEP^YAP5SA^ tissues with unengineered FB^WT^/HEP^WT^ controls, and found that engineered tissues only proliferated more than controls when treated with DOX for 7 days (Fig. S3G). We were curious whether a short pulse of DOX would also be sufficient to kickstart the HEPs into a proliferative regime, but found that FB^HTWR^ /HEP^YAP5SA^ tissues required sustained DOX treatment to reach a maximal proliferative effect at 7 days, although some effect was observed with shorter 3 and 5 day treatments (Fig. S3H). We explored the kinetics of this proliferative response and found that HEPs reach maximal proliferation 5 days after adding DOX, which aligns well both with the duration of DOX required for proliferation and the kinetics of YAP target gene activation we found previously (Fig. 2K). We also tested whether HEPs would revert back to quiescence upon withdrawal of DOX, a response critical for clinical safety. Via Ki67 imaging we found that proliferation drops after removal of DOX and returns to baseline within 5 days of DOX withdrawal (Fig. 2L). These proliferation kinetics were mirrored by engineered cell number changes: luminescent signal increased during the first 7 days of the experiment during DOX treatment, then plateaued during the subsequent 7 days when DOX was withdrawn (Fig. 2M). Together, these experiments demonstrate our success in designing a synthetic biology toolkit that enables on-demand expansion of 3D engineered liver tissue.

### YAP/GF expanded HEPs retain their differentiated cell state and exhibit a tradeoff between function and proliferation

Prior to implantation into a host, we felt that it was critical to assess the changes that occur within HEPs when they are exposed to combined YAP/GF stimuli. Proliferation is classically associated with a reduction in cellular functions, as cells need to prioritize energy expenditure(*41*). Furthermore, some HEP expansion strategies currently employed in the field result in clinically undesirable dedifferentiation of HEPs into a progenitor, bipotential, or fetal-like cell state(*34*, *36*). To probe the effects of synYAP/GF on the proliferative, functional, and differentiation status of HEPs we performed single nucleus RNA sequencing (snRNA-seq) on engineered 3D liver tissues containing WT or HTWR FBs and WT or YAP5SA HEPs after 1 week of doxycycline induction. Five replicates were assayed per condition, yielding a total data set after doublet removal and QC of 18,991 nuclei, of which 4615 (24%), 6962 (37%), and 7414 (39%) were of HEP, EC, and FB, respectively (Fig. 3A, S4A). Sequencing quality was consistent across batches, conditions, and cell type, and comparable to previously reported snRNA-seq QC metrics (Fig. S4B,C,D).

**Figure 3.**
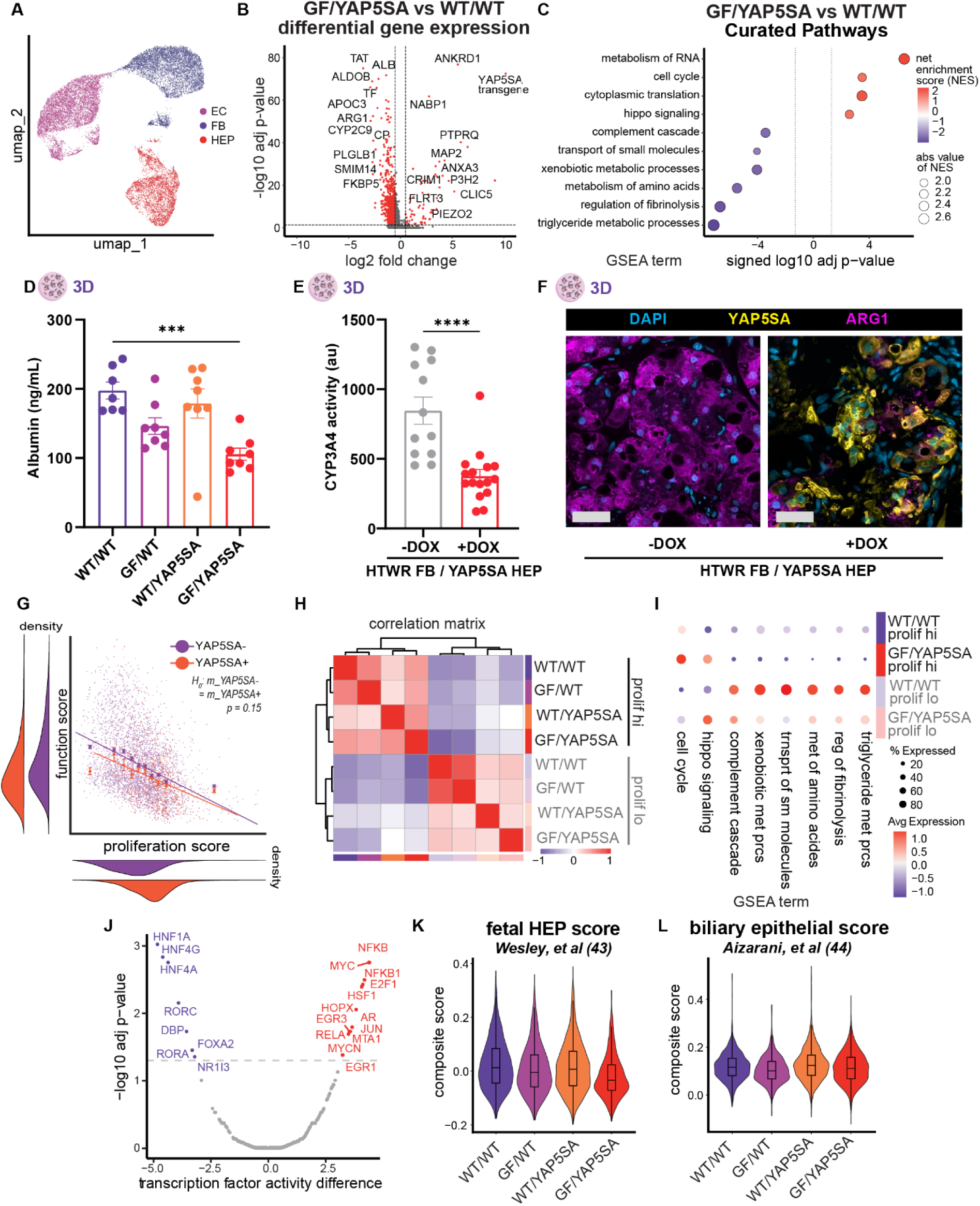
YAP/GF expanded hepatocytes retain their differentiated cell state and exhibit a tradeoff between function and proliferation. **(A)** UMAP clustering of complete data set revealing distinct EC, HEP, and FB clusters. **(B)** Differential gene expression HEPs in FB^HTWR^ /HEP^YAP5SA^ vs FB^WT^/HEP^WT^ samples. **(C)** Pathway analysis of genes differentially expressed by HEPs in in FB^HTWR^ /HEP^YAP5SA^ vs FB^WT^/HEP^WT^ samples. **(D)** Measured albumin protein production of 3D liver tissues after 7 days of synGF/YAP5SA expression. **(E)** Measured CYP3A4 enzyme activity of 3D liver tissues after 7 days of synGF/YAP5SA expression. **(F)** Immunohistochemical protein staining of arginase 1 of 3D liver tissues after 7 days of synGF/YAP5SA expression. Scale bars 50 µm. **(G)** Proliferation vs function module score of all HEPs data set labeled by YAP5SA transgene positivity. Best fit lines of YAP5SA +/- lines compared via ANOVA. **(H)** Hierarchical clustering of HEP from each experimental conditioned binned by proliferation score (hi – 75^th^ percentile or lo 25^th^ percentile). **(I)** Pathway module scores by HEPs in FB^HTWR^ /HEP^YAP5SA^ vs FB^WT^/HEP^WT^ samples binned by proliferation score. **(J)** Inferred transcription factor activity of FB^HTWR^ /HEP^YAP5SA^ HEPS compared to FB^WT^/HEP^WT^ HEPs. **(K)** Fetal HEP composite scoring by experimental condition. **(L)** Biliary epithelial composite scoring by experimental condition. (p *<.05, ** < .01, *** < .001, **** < .0001; D: one-way ANOVA, E: t-test)

We first assayed changes in the stromal cell populations of our liver implants, and found that YAP5SA expression in HEPs did not cause any detectable changes to the ECs and FBs cocultured in those samples (Fig. S5B,C). Further, GF expression by the FBs caused only modest changes to the EC and FB transcriptomes, again suggesting that these cells are not substantially altered by the process of expressing the HEP mitogens (Fig. S5B,C). On the other hand, we detected many significantly differentially-expressed genes between HEPs in different YAP and GF treatment groups, especially those exposed to both YAP and GF activation (Fig. 3B, S5A). HEPs in FB^HTWR^ /HEP^YAP5SA^ more strongly expressed genes associated with cell cycle, RNA metabolism, and translation compared to engineered FB^WT^/HEP^WT^ controls, consistent with increased HEP proliferation we observed with dual YAP/GF activation (Fig. 3C, S6A). On average, HEPs in FB^HTWR^ /HEP^YAP5SA^ samples concurrently downregulated a variety of HEP-specific functional processes including peptide, lipid, and xenobiotic metabolism, small molecule transport, and production of clotting factors and complement. Other HEP functions were not significantly altered, but bile acid transport and urea cycle functions displayed decreasing trends, and HEPs appeared to be shifting towards an increased iron and glycogen storage phenotype (Fig. S6B). While YAP and GF activation alone each only slightly influenced these pathways, combined YAP and GF stimuli cooperated to produce the strongest effects on proliferation and function, consistent with the synergistic relationship we had observed phenotypically (Fig. 2H,I, Fig. S7A,B,C,D).

To validate these findings, we assayed three critical aspects of HEP function – albumin production, drug metabolism, and urea cycle activity – in our engineered liver tissues *in vitro*. Consistent with our sequencing data, both albumin production and cytochrome-p450 activity were reduced and urea cycle enzymes like arginase-1 were downregulated (Fig. 3D,E,F).

Given only a subset of HEPs in our engineered liver tissues proliferate at any given time, we were curious if the observed reduction in function of FB^HTWR^ /HEP^YAP5SA^ samples is driven by a decrease in function across all HEPs in that sample, or only by those that are cycling. To address this question, we calculated a proliferation and a function score for each cell in our sequencing dataset and found that proliferation and function were anticorrelated: individual cells that were more proliferative tended to be less functional, and vice versa (Fig. 3G). Interestingly, this anticorrelation existed even in wildtype HEPs and expression of YAP5SA did not significantly alter this relationship, but instead drove an overall cell population shift towards high proliferation and low function scores. We also found that function genes more strongly anticorrelated to proliferation genes than to Hippo signaling genes, further supporting that proliferative status, rather than YAP5SA transgene expression, is more associated with HEP function (Fig. S8A,B). To further assess this possibility, we binned HEPs across the entire data set by proliferative score and found that HEPs hierarchically clustered by proliferative state rather than experimental condition (YAP5SA or GF induction) (Fig. 3H, Fig. S8C,D). YAP5SA+ HEPs from FB^HTWR^ /HEP^YAP5SA^ samples with low proliferation scores retained functional capabilities similar to WT HEPs with low proliferation scores, despite their elevated Hippo signaling activity (Fig. 3I, Fig. S8E). Moreover, both proliferating HEPs in FB^HTWR^ /HEP^YAP5SA^ and FB^WT^/HEP^WT^ downregulate HEP specific functions in a similar fashion. We further confirmed this relationship histologically: proliferating HEPs in both uninduced and YAP/GF activated tissues downregulate production of albumin and arginase-1 compared to neighboring non-proliferating HEPs (Fig. S9A,B).

To characterize the functional state of HEPs after halting proliferative stimulation, we compared YAP/GF engineered tissues that were never treated with DOX, treated constantly for 14 days, or treated for only the first 7 days followed by a 7 day recovery period without DOX induction. We assayed albumin and urea production and CYP enzyme activity and found that while tissues dosed with DOX for 7 days function better than those treated for 14 days, our 7 day treated samples were not more functional than unexpanded controls, despite the proliferation-induced increase in number of HEPs per graft (Fig. S10A,B,C,D,E). Further interrogation of these samples with bulk RNA sequencing showed HEPs cluster by DOX treatment duration, and that on a per-cell basis the functionality of HEPs, based on HEP-specific gene expression profiles, were similar following the 7-day DOX treatment period as compared to after a week of recovery without DOX (Fig. S11A,B,C,D,E). This suggests that once expanded, withdrawal of DOX does not induce HEPs in our system to spontaneously recover function but does successfully prevent additional function loss.

Mechanistically, we were interested in what might ultimately be driving the observed changes in HEP function and proliferation. Using decoupleR we assayed for predicted transcription factor activity, and found a variety of transcription factors known to regulate both proliferation and HEP function to be inversely perturbed upon YAP and GF activation, some of which were altered at the transcript level (Fig. 3J, Fig. S12A)(*42*).

Beyond reduction in function, dedifferentiation of HEPs to a fetal, bipotential or biliary cell fate has also been reported in some expansion strategies and is translationally undesirable. In response to synGF and YAP5SA expression we did not see emergence of classical fetal HEP or biliary markers such as AFP, CYP3A7, IGF2, KRT19 or SOX9, while mature HEP markers were comparatively maintained (Fig. S12B). We also scored our samples on curated gene sets from fetal human liver(*43*) and mature biliary epithelium(*44*) and did not see emergence of a fetal or biliary phenotype (Fig. 3K,L). Together, these findings support that YAP and GF activation can drive an enhanced proliferative effect in 3D engineered hepatic tissues without inducing an undesirable fetal, bipotential, or biliary cell fate.

### Synthetic control over YAP and GFs licenses proliferation of implanted ectopic liver tissues in situ

Given our promising *in vitro* results, we were eager to investigate if control over GFs or YAP signaling could modulate the proliferation of engineered liver tissues post implantation, a critical first step towards *in situ* expansion. To test whether GFs alone would be sufficient to induce proliferation of implanted HEPs in an *in vivo* setting, we first implanted engineered liver tissues containing WT HEPs and GF producing FBs into healthy NSG mice (Fig. 4A). Mice were fed DOX chow to induce GF expression and injected daily with EdU to label proliferating cells. After 1 week of GF induction, implants were retrieved and assessed histologically. Compared to a pre-induction baseline, we observed a 50% increase in serum human albumin in implants with HGF and *TGFα* expressing FBs (Fig. S13A). We further observed a modest increase in proliferation of HEPs exposed to HGF and *TGFα* via longitudinal EdU labeling (Fig. 4B, S13B). To assess whether the effects of synthetically produced GFs were isolated locally, as desired, or providing systemic stimulation to distant tissues, we assessed the host livers histologically and measured mouse liver to body weight ratio and found no changes in response to synGF production (Fig. 4C, Fig. S13C,D).

**Figure 4.**
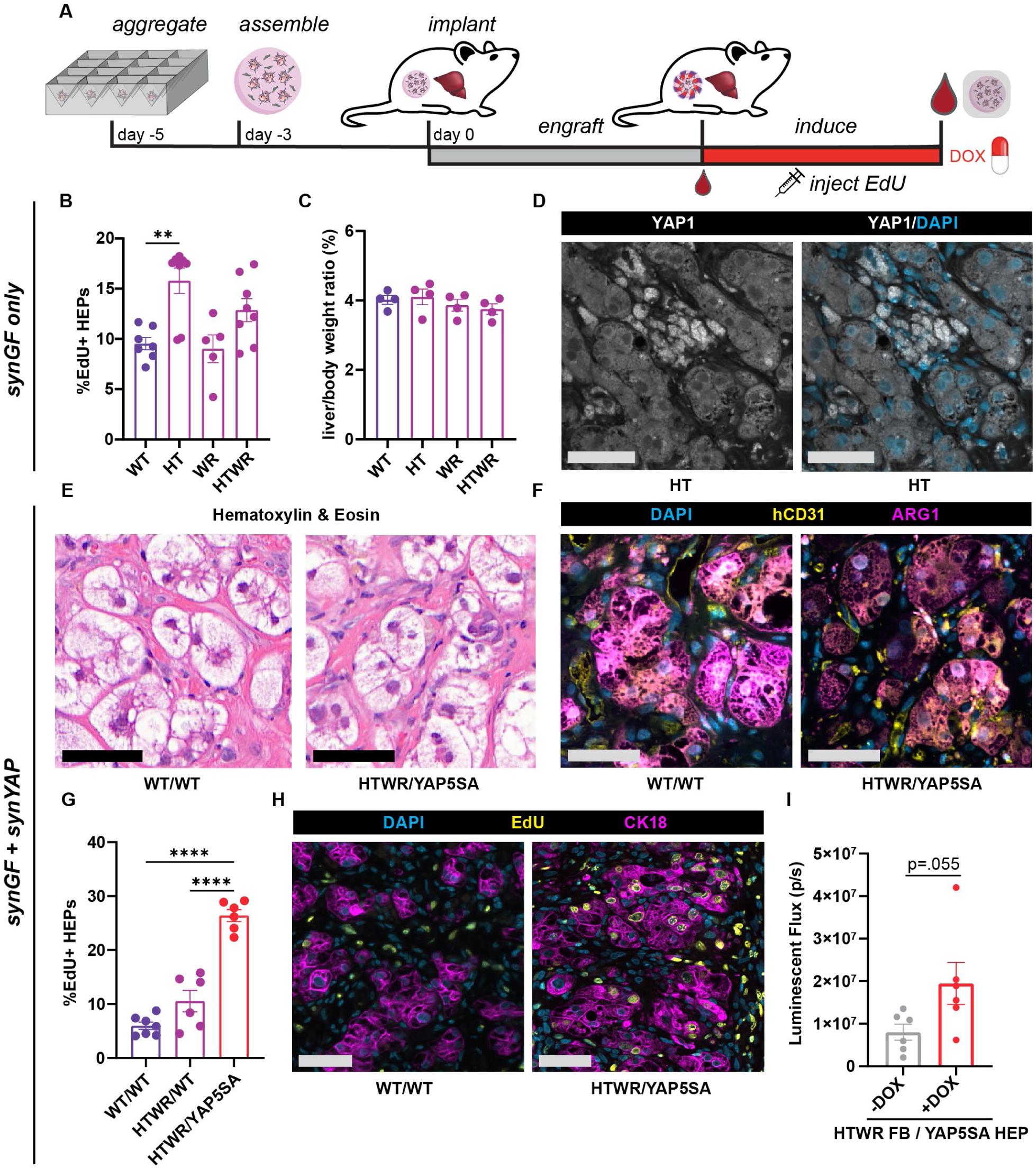
Synthetic control over YAP and GFs licenses proliferation of implanted ectopic liver grafts in situ. **(A)** Schematic of ectopic liver graft assembly and implantation. **(B)** Quantification of HEP proliferation during 1 week *in vivo* induction of synGFs in implanted liver tissues (N:8,n:2). **(C)** Liver to body weight ratio in mice after 1 week of *in vivo* induction of synGFs (N:4, n:2). **(D)** Immunohistochemical staining of YAP1 in synHT expressing ectopic liver tissues. Scale bars 50 µm. **(E)** H&E staining of FB^WT^ /HEP^WT^ and FB^HTWR^ /HEP^YAP5SA^ ectopic liver tissues after 1 week of DOX induction. Scale bars 50 µm. **(F)** Immunohistochemical staining for human blood vessels (hCD31) and human HEPs (Arg1) of FB^WT^ /HEP^WT^ and FB^HTWR^ /HEP^YAP5SA^ ectopic liver tissues after 1 week of DOX induction. Scale bars 50 µm. **(G)** Quantification of proliferation of FB^WT^ /HEP^WT^, FB^HTWR^ /HEP^WT^, or FB^HTWR^ /HEP^YAP5SA^ liver tissues during 1 week of DOX induction (N:3,n:2). **(H)** Immunohistochemical staining of proliferating HEPs (labeled by HEP marker CK18) from samples quantified in G. Scale bars 50 µm. **(I)** Luminescence of implanted FB^HTWR^/HEP^YAP5SA^ tissues measured via intravital imaging after 1 week of DOX induction compared to uninduced controls (N:6,n:1). Replication reporting: N: number of animals, n: number of implants per animal (p *<.05, ** < .01, **** < .0001; B,C,G: one-way ANOVA, H: t-test; B: data shown from one representative experiment, see supplement for additional repeats.)

In light of the modest increases in proliferation observed with synGFs alone, we suspected that implanted HEPs may be constrained by a density-related checkpoint, as seen in our cultures *in vitro*. We stained for YAP in both FB^WT^ and FB^HT^ tissues and found that YAP was excluded from the nuclear regardless of GF treatment (Fig. 4D, S14A), suggesting that synYAP expression may be beneficial for enhancing proliferation of implanted HEPs, similar to our findings in cultured constructs. To test this hypothesis, we implanted FB^WT^ /HEP^WT^, FB^HTWR^ /HEP^WT^, and FB^HTWR^ /HEP^YAP5SA^ tissues and induced GF and YAP activity for 1 week with DOX. Dual engineered FB^HTWR^ /HEP^YAP5SA^ implants were well tolerated by their hosts: we observed no loss of body weight or changes in animal behavior upon induction of GFs and YAP. Implants histologically also appeared similar: there was no evidence of increased fibrosis, immune infiltrate or nodules, and HEPs morphologically appeared similar on H&E across conditions and when compared to ectopic liver reported elsewhere (*31*, *45*) (Fig. 4E, S15A,B,C). FB^HTWR^ /HEP^YAP5SA^ implants were also well vascularized without need for advanced vascular patterning strategies (Fig. 4F). Consistent with our *in vitro* results and sequencing data, synthetic expression of GFs resulted in a reduction in serum human albumin concentration, which was further exacerbated by induction of YAP5SA in HEPs (Fig. S16A). Notably, however, we found that while GF expression alone once again resulted in only modest increases in proliferation, FB^HTWR^ /HEP^YAP5SA^ dual-engineered implants exhibit a striking 500% increase in EdU incorporation compared to unengineered FB^WT^ /HEP^WT^ tissues, but do not form tumors (Fig. 4G,H S17A,B,C,D). Using intravital imaging, we further show that this proliferative effect results in a doubling of engineered HEPs after just one week of combined YAP/GF stimulus compared to uninduced controls (Fig. 4I, Fig. S16B). Together these data define our combined synGF/YAP strategy as a promising approach to drive on demand growth of ectopic liver tissues *in situ* post implantation, and further serve as a proof of concept that BOOSTing therapies towards therapeutic scale could indeed be possible.

## Discussion

In this study we define the first steps towards an unconventional approach to cell therapy scale up: engineering a small construct and then inducing it to grow *in situ*. This strategy, which we have named BOOST (<u>B</u>ioengineered <u>O</u>n-demand <u>O</u>utgrowth via <u>S</u>ynthetic biology <u>T</u>riggering), could provide several key advantages, including circumventing the need for a large pool of cellular raw materials and bypassing the formidable challenge of generating a rapidly perfusable construct that can survive the engraftment period. We leveraged a set of 2D and 3D human liver culture models to define the pairing of YAP and GF signals as a set of sufficient cues to stimulate proliferation, but that also do not lead loss of human HEP cell fate, even in dense 3D engineered tissues. We then designed a toolkit to synthetically modulate these cues locally within an engineered liver tissue and showed that combined synthetic manipulation of YAP and GF enabled on-demand expansion of human hepatic tissue *in vitro* and *in vivo*. As such, this work serves as an exciting proof-of-concept demonstration that scale up of tissues via growth could be possible.

Beyond the technical advances presented herein, we believe that our system, as a fully human engineered 3D liver tissue in an implant setting, has provided unique and valuable insight into the nuances and drivers of human HEP proliferation. Indeed, although nearly a century of literature exists on the study of liver regeneration, the vast majority is based on rodent liver regeneration, where overexpression of HGF or TGFα, or constitutive activation of YAP signaling (via genetic manipulations or drugs) each alone are sufficient to trigger liver hypertrophy (*46–49*). In contrast, our gain-of-function studies demonstrate that expression of multiple growth factors and YAP5SA are simultaneously required to trigger human HEP proliferation *in vivo*. While these findings are consistent with the requirement for concurrent activation of multiple pathways to trigger *ex vivo* human HEP expansion(*33*, *34*, *36*, *50*) and with the role for YAP signaling in liver regeneration (*51*), the mechanism by which YAP and GF signaling synergize to trigger human HEP proliferation is yet unclear. It is known that YAP integrates mechanical information from to cell-cell and cell-ECM contacts, polarization, and cytoskeletal tension to induce contact inhibition of proliferation in epithelial cell sheets(*52–54*). However, while YAP can be activated upon GF treatment and is sufficient to drive proliferation in some cell types(*39*, *55*, *56*), our findings suggest that this is not the case in human HEPs. Instead, YAP and GF signaling appear to be integrated orthogonally through a functional “AND” gate. The molecular underpinnings of such a multi-factorial regulatory checkpoint system for growth control in human HEPs, however, remains unclear and warrants future study. As such, our work highlights key differences between human and rodent HEP proliferation regulation and underscores the need for more experimental models and studies focused on human liver biology.

While here we offer an approach to overcoming a critical challenge in tissue engineering –3D tissue scale-up – our studies align well with the overall momentum in the mammalian synthetic biology and cell therapy fields to control therapeutic behavior after implantation(*26*, *57*). Although these synthetic control modules have predominately been applied to immunologic applications, development of therapies such as CAR-T cells provide a roadmap to clinical translation. Now, use of synthetic biology has been expanded to address challenges unique to tissue-based therapy, such as vascularization(*58*) or tissue morphogenesis(*59*). Here we instead leverage the power of synthetic biology to define an approach to synthetically trigger proliferation of an implanted tissue for the purposes of *in situ* scale up: BOOST. While induction of proliferation for endogenous tissue regeneration *in vivo* has precedents(*17*, *19–21*), as does stimulation of immunologic cell therapy product proliferation *in vivo* post administration(*26*, *60–62*), BOOST integrates ideas from synthetic biology and regenerative medicine into tissue engineering, approaching the challenge of solid tissue therapy scale up from an orthogonal, genetic approach to conventional fabrication focused scaling methods. Realization of this approach required not only the definition of a set of cues that could trigger 3D liver tissue proliferation, but also the genetic engineering of primary human HEPs to control these cues, a notoriously challenging cell type to manipulate. Whereas previous reports have relied on using artificially induced cycles of hepatic injury for HEP engraftment and expansion(*31*, *63*, *64*), here we successfully relieved this dependence and achieved control over implant growth in healthy mice. While growth control in any cell or gene therapy application raises concerns of possible tumorigenicity, we validated that synthetic YAP/GF system is switchable and does not result in the formation of tumors at the implant site or in distant organs, even when constantly induced (Fig. S13, S17). This is likely because the endogenous tumor suppressor machinery in the engineered HEPs remains intact and synthetic expression of GFs is confined to the implant site alone. Moreover, liver tissues engineered with BOOST could also be equipped with a kill-switch to shield against adverse events (*25*).

While control of tissue growth is a critical first step towards translation of BOOST, during these studies we observed that induction of proliferation led to a reduction in function of expanded HEPs. This functional reduction appears to result from a natural tradeoff between function and proliferation conserved across many mammalian cell types(*65–68*). We expect that an additional set of “maturation” factors, however, are likely needed to actively recover function, as has been ubiquitously required in *in vitro* HEP expansion strategies(*33*, *35*, *37*, *69*, *70*). An exciting next step for BOOST towards clinical translation will be to expand on our existing genetic circuitry to not only control HEP proliferation but also to enhance HEP function.

The work presented herein serves as an exciting proof of concept demonstration of on-demand implanted solid cell therapy expansion and defines the first steps towards clinical translation of BOOST. We anticipate that as additional regulators of human cell proliferation and function inevitably emerge, such insights will only further facilitate more efficacious implementation of this strategy in the years to come. We envision that BOOST could be useful to many other solid cell therapies, such as pancreatic or cardiac tissue, that are also currently similarly constrained by scale-up associated challenges. Together, this work helps lay the foundations for a future of “smart” solid organ cell therapies that can be scaled to a patient’s needs, and thereby offer treatment for numerous, previously incurable, diseases.

## Materials and Methods

### Cloning and lentiviral production

Starting plasmids were obtained through addgene or generated via gene block synthesis. For all house made constructs, an all in one TET system was used as the primary backbone. Genes of interest were cloned into the first molecular cloning site using Gibson assembly, and the 2A sequence was excised. To make the integrated luciferase reporter, the PuroR sequence was excised and replaced with a firefly luciferase. Completed plasmids were sequenced and used to produce lentivirus via conventional methods as described previously(*24*). Supernatant containing lentivirus was concentrated using lentiviral concentration solution, aliquoted and stored at -80C until use.

### 2D HEP culture

HEPs were plated on collagen I (100 ug/mL) coated 96 well plastic plates in 150 uL/well of Advanced DMEM + 1x B27-vit A + P/S (Adv DMEM++). 10,000 primary human HEPs (Gibco donor 8339), or two consecutive rounds of 50,000 PHHs, were used to seed sparse and dense cultures, respectively. After 1 hour of incubation at 37C to allow for cell attachment, cultures were washed gently to remove non-adherent cells and fresh media added.

After 24 hours, cells were switched into Adv DMEM++ supplemented with test compounds (see supplemental materials list) or conditioned media (10x concentrated, reconstituted to 1x in adv DMEM++) and EdU (5uM, when relevant). After 48 hours, cells were fixed for 20 minutes with 4% paraformaldehyde or lysed for PCR or western blot.

### 3D liver graft assembly and culture

On the day of aggregate assembly, neonatal normal human dermal fibroblasts (nNHDFs) were growth arrested with 10 ug/mL mitomycin C for 2.5 hours. Arrested nHDFs were washed 5 times with media prior to use. Murine 3T3-J2 fibroblasts were used for the experiment in Figure 2G to enable species separation of bulk RNA analysis by qPCR of aggregate cultures.

HEPs were thawed and directly combined with FBs and seeded into pluronics F-127 passivated house-made pyramidal aggrewell molds at a density of 5M:1M HEPs:nNHDF per well in a 1:1 mixture of EGM2/ITS medias. Cultures were spun down at 60g for 6 minutes and carefully transferred to an incubator for 2 days.

2 days following aggregation, aggregates were collected and assembled into liver grafts. Briefly, PDMS gaskets (4mm/ 6mm ID, 8/10 mm OD (*in vitro* 25 ul grafts/ *in vivo* 50 uL grafts)) were punched, sterilized, and placed into dry, pluronics treated non TC well plates. Harvested aggregates were resuspended in fibrinogen (20 mg/mL) with 0.25 U/mL human thrombin and human umbilical endothelial cell (HUVEC) suspension and pipetted into PDMS gaskets. Final grafts contained 125k/125k/250k of PHH/HUVEC/nNHDF per 25 uL of 10 mg/mL fibrin hydrogel. After 1 hour of gelation at 37C, grafts were fed with EGM2/ITS (unless otherwise noted) and detached from the bottom of the culture dish. Media was changed every 1-2 days.

For grafts requiring lentiviral infection, PHHs were thawed into ITS media with 8 ug/mL polybrene and 50 MOI tet-YAP5SA lentivirus, plated into passivated aggrewell molds, and spinoculated at 2000 rpm for 45 minutes. After overnight incubation, HEPs were collected, washed 5 times, and then incorporated into aggregates as described above.

Luminescent grafts were formed in white clear bottom well plates and imaged at relevant time points 20 mins after addition of 150ug/mL D-Luciferin. RLU reported as the luminescent flux normalized by day 0 graft initial signal (by graft) and average uninduced graft signal (by time point).

All primary human cell lines were used between passages 5 and 7.

### In vivo implantation and assessment of ectopic liver grafts

All procedures were approved and overseen by the MIT committee on animal care. 3 days after graft assembly, 50 uL (6mm) grafts were implanted into 8-10 week old female NSG mice (Jackson, 005557). Briefly, mice were anesthetized with isoflurane, abdomen clipped and sterilely prepared, and incisions made through the skin and abdominal wall medial to the 4^th^ nipple. Implant was sutured using 6-0 nylon suture to an exteriorized piece of the perigonadal fat pad. Incisions were closed with 5-0 nylon suture (abdominal wall) and wound clips (skin). Bilateral implantations were performed unless otherwise noted. Animals were administered Buprenorphine SR or Ethiqa XR on the day of surgery for pain management.

To induce TET genetic circuits *in vivo* animals were fed *ad libitum* with doxycycline chow (625 ppm). Animals were initiated on the doxycycline chow two days prior to implantation and maintained on this diet for the duration of the experiment.

To monitor implant function, 100 uL of blood was collected from the saphenous vein weekly (alternating legs) into heparinized microcapillary tubes, spun at 1500xg for 5 minutes, and serum frozen at -20C for later analysis.

To label implant proliferation, mice were injected intraperitoneally once daily for 1 week with 50 mg/kg EdU (Carbosynth) in sterile saline.

To monitor luminescent signal of implanted HEPs, animals were subcutaneously injected with D-Luciferin at a dose of 150 mg/kg. 20 minutes after injection, animals were imaged on a Perkin Elmer IVIS Spectrum system to detect luminescent signal. Signal was integrated within an ROI drawn around each animal’s abdominal cavity and reported as total luminescent flux. Animals were euthanized via cervical dislocation under deep anesthesia, implants retrieved and fixed, and terminal blood draws collected via retroorbital bleed. When applicable, animal liver to body weight ratio was also assessed at end point.

### Histological assessment of engineered liver grafts

Engineered liver tissue explants or *in vitro* cultured liver grafts were fixed over night at 4C with 10% neutral buffered formalin and paraffin processed and embedded. Tissues were sectioned at 5um thickness and immunofluorescent stained with relevant chemistries and protocols. Briefly, slides were deparaffinized, steamed in sodium citrate antigen retrieval buffer, and blocked/permeabilized in a mixture of .05% Triton, 5% normal donkey serum in PBS. Following blocking and permeabilization, EdU detection was performed using a house made mixture of water, 10X TBS, 10mM CuSO4, 1 M sodium ascorbate, and 7 ng/mL fluorescent azide (click chemistry tools). Primary antibodies (see supplemental materials list) were incubated overnight in a humidity chamber at 4C, secondary antibodies for 1 hour at room temperature. Slides were mounted with prolong Gold Antifade Mountant.

Stained slides were imaged on a 3D HisTECH Panoramic Scanner 2000 or TissueFAXS SL Fluorescent Slide Scanner. Resulting images were imported into QuPath for quantification.

H&E staining was performed using an automated staining workstation. Masson’s modified trichrome staining was performed in house. Resulting slides were imaged via slide scanner.

### Image analysis and quantification

Images of 2D stained cultures were imported into Cell Profiler (v4.2.6) to quantify the number of cells present per high powered frame, the percent Edu+ PHHs, and YAP or YAP5SA nuclear/ cytoplasmic ratio. Nuclei were detected using DAPI staining and filtered for live HEPs based of nuclear morphology (bright, very small nuclei were excluded as apoptotic). To quantify EdU positive cells, a secondary cell detection was performed on the EdU channel, and DAPI+ nuclei filtered by this detection were denoted Edu positive. Percent Edu+ HEPs was calculated by dividing the total number of EdU positive cells detected in 15 FOV per well by the total number of cells detected. To quantify YAP nuclear to cytoplasmic ratio (NCR), nuclei were identified as before, and a 10 pixel annulus was established around each nucleus. YAP NCR was calculated as the YAP staining intensity in the nucleus divided by that of the annulus. For ease of interpretation, in some panels YAP NCR was binned into primarily nuclear (N, NCR > 1.1), primarily cytoplasmic (C, NCR < 0.9), or both nuclear and cytoplasmic (N/C, 0.9 < NCR < 1.1) as reported previously(*52*).

Images of stained 3D liver graft sections were imported into QuPath (v0.5.0) and used to quantify the number of EdU/Ki67+ HEPs. HEPs were first identified using a pixel classifier on the CK18 channel of the image. HEP annotations were then eroded by 3 µm to remove any HEP-adjacent FBs. Nuclear detection was then performed on each HEP annotation using DAPI staining. Nuclei were then classified as label (eg EdU, Ki67) positive using a custom script. Nuceli were considered positive if the nuclear staining intensity to be above a threshold and more than 2-fold higher than both the cytoplasmic staining intensity and the average cellular intensity of all HEP detections. This data is reported as %EdU or Ki67+ HEPs, where one data point represents at least 500 HEPs from at least 2 different planes of section of one liver graft. Absolute quantification of HEP area and HEP nuclei in Supplementary Figure X were performed as described above using slides from 50um step sections through the entirety of each liver tissue.

### CYP enzyme activity

CYP3A4 activity was assayed using the Promega P450 Glo assay as described per manufacturer’s instructions. Briefly, liver grafts were first treated with CYP3A4 activating small molecule rifampicin (25 uM) for 2 days prior to the assay. Grafts were then incubated with luciferin-IPA probe for 1 hour in DMEM P/S. After incubation, samples of media were developed for 20 minutes in duplicate and luminescence measured on a plate reader.

### ELISA

Albumin was detected using a conventional sandwich ELISA protocol. Briefly, 96 well assay plates were coated with an anti-albumin antibody in a basic sodium carbonate/bicarbonate coating buffer, blocked and incubated with an eight point standard curve and sample for 1 hour. Plates were washed, incubated with Albumin-HRP secondary antibody for 1 hour, washed and developed with Ulta-1 step TMB. Development was quenched with 0.5N HCl after 4 minutes and absorbance measured on a plate reader.

ELISA for detection of TGFα and HGF was performed as described above using kits from R&D systems with the addition of a streptavidin secondary conjugation to enhance detection.

### synGF conditioned media and WNT reporter assay

GF producing NHDFs were grown to confluence in 10 cm dishes and then treated with 1 ug/mL doxycycline for 4 days. After conditioning period, media was harvested, centrifuged to remove cell debris, and passed through a 0.2 µm filter to sterilize. Aliquots of conditioned media were frozen at -80 C and were not freeze thawed more than once.

A HEK reporter line was generated expressing the pBARLS Wnt reporter(*71*). To assay WNT activity of conditioned medias, 100k BARLS HEK were plated per well of a 96 well plate in NHDF conditioned media diluted 2 parts to 1 part with HEK growth media. After 24 hours incubation, media was removed and samples lysed with glo-luciferase reagent and luminescence quantified using a plate reader. Fold change reporter activity is reported relative to WT conditioned media.

### Western Blotting

Cells were lysed on ice with RIPA buffer and normalized using a Bradford total protein assay. Samples then were diluted with sample buffer and 5% BME, thermally denatured, and run on 4-12% Bis Tris Gels using MOPS running buffer. Gels were transferred for 2 hours at 180 mA and blocked with 5% BSA or milk in TBST. Primary antibodies (see supplemental materials list) were incubated at 4C overnight, secondary antibodies for 1 hour at room temperature. Blots were developed using Super Signal Pico Plus chemiluminescent substrate and imaged on an iBright gel imager. For presentation, blots were contrast adjusted and background subtracted.

For detection of MET and EGFR activation, HEPs seeded at a density of 300k/ 6 well were treated with NHDF conditioned media for 30 mins. 2.5 minutes prior to lysis, pervanadate solution (100 mM sodium orthovanadate in PBS + 0.25% hydrogen peroxide) was directly spiked into the media to reduce phosphatase activity. Phosphorylated proteins were first detected, then membrane stripped and restained for pan protein. Quantifications were performed using ImageJ on raw blot images, normalized to relevant loading controls.

### RNA isolation, qPCR, and bulk RNA sequencing

RNA was isolated using phenol chloroform extraction and purified using Quiagen RNAeasy mini or MinElute RNA cleanup kits. First strand cDNA synthesis was performed using qScript cDNA SuperMix. qPCR was performed in duplicate in a 384 well plate format on a BIORAD 384 well themocycler. Primers were used at a concentration of 1.25uM were designed to bind human templates only (see supplemental materials list); species specificity was validated via NCBI nucleotide blast and confirmed experimentally.

For bulk RNA sequencing, 4 replicates per condition of whole *in vitro* liver tissues were digested using a Quiagen TissueLyser II and RNA extracted using Quiagen RNAeasy mini kits. Libraries were generated after RNA QC (all samples RIN > 9.9) with poly A based NEB UltraII RNAseq and sequenced on 4 lanes of 50 PE F3 Singular flowcell with a Singular G4 sequencer. All samples surpassed standard QC metrics for bulk RNA sequencing. Data were processed with DESeq2, normalizing by a panel of housekeeping genes determined to have non-varying expression across cells and conditions from snRNA-seq data. All data presented queries only HEP specific genes (those with ≥ 10 fold higher expression in HEP containing samples compared to HUVEC/FB only controls).

### Single nucleus RNA sequencing

*In vitro* 3D liver tissues were assembled as described, and snap frozen in liquid nitrogen prior to analysis. Nuclei were isolated using a previously reported protocol. Briefly, grafts were dissociated by a Quiagen TissueLyser II, filtered, and flow sorted using a SONY FACS sorter. Single nuclear sequencing libraries were generated using the ddSEQ Single-Cell 3’ RNA-Seq Kit from BIORAD. Samples were sequenced on a NextSeq 2000 with a P3 100 cycle kit (54 bp read 1, 8 bp index 1, 8 bp index 2, 68 bp read 2). Omnition analysis software (v1.1.0) was used to align sequencing reads to a custom reference genome, consisting of hg38 appended with custom transcripts for transgene regions unique to engineered cells. Seurat objects outputted by the Omnition pipeline were imported and merged into a single object in RStudio (RStudio 2024.12.1). Standard Seurat (v5.2.1) pre-processing was performed, including log normalization, PCA analysis, UMAP clustering (using unbiased resolution and dimensionality reduction). Populations containing doublets or low QC metrics (low UMI or RNA feature count, and/or high mitochondrial transcript count) were manually excluded from the dataset. Subsets of HEPs, FBs, and ECs were clustered independently to explore variation within cell type populations. Differential gene expression was computed between conditions of the same cell type. GSEA was performed using the complete gene lists from MSIGDBr GO:BP and CP:REACTOME. To compare gene set expression across conditions, HEPs were scored on leading edge genes of GSEA form the comparison between FB^WT^/HEP^WT^ and FB^HTWR^ /HEP^YAP5SA^ HEPs. Proliferation and function scores were calculated by module scoring on REACTOME:CELL CYCLE and the geneset emerging from the intersection of 3 HEP identity gene sets from healthy human HEPs (C8:Cell Type Signature:AIZARANI_LIVER, clusters 11, 14, and 17), respectively. Anova was used to compare the linear regressions of YAP5SA transgene positive and negative subpopulations. Linear modelling was performed using the stats:lm() function on scaled (stdev 1, mean 0) function, identity, or GOBP_HIPPO_SIGNALLING module scores. HEPs were considered proliferation score hi or lo if their cell cycle score was in the top, or bottom quartile of the combined HEP data set, respectively. Hierarchical clustering was performed on correlations of aggregate gene expression of cells subclassified by proliferation score bin (hi, lo) and condition. Module scores for fetal HEPs and cholangiocytes were calculated based on a curated gene set of marker genes from Wesley et al (*43*) and C8:Cell Type Signature: AIZARANI_LIVER_C4/ C7_EPCAM_POS_alBILE_DUCT_CELLS, respectively. Inferred transcription factor activity was computed using decoupleR on psueodbulk average gene expression by sample(*42*).

### Statistics and reproducibility

Experimental data were analyzed and processed in MATLAB and Microsoft Excel. Statistical tests and plotting were performed in GraphPad Prism software (v10). When applicable, tests for normality were performed to determine appropriate statistical methods. No data was excluded from the analyses. For some experiments, only a single experimental repeat was presented in the main figure for improved clarity. Additional experimental repeats can be found in the supplemental figures referenced in the text, wherein filled symbols denote the repeat shown in the main figure, and different shapes denote different experimental repeats. Some data from the main figures is reproduced in the supplemental figures for ease of comparison. Replication reporting is included in figure captions. Experiments were not randomized or blinded.

## Acknowledgments

We would like to thank H.H.G. Song, L. Perry, K.A. Gagnon, J.M. Bays, M. Uroz, J. Eyckmans, S. Bakre, G. Gini, A.X. Chen, S. March, K. Grzelak, A. Westerfield, C. Martin-Alonso, C. Misturado, and H. Fleming for thoughtful discussion and enabling scientific training, and L. Kelleher, A. Michaels, L. Ch’ng, S. Kangiser, and B.F. Bard for administrative and technical support. We would like to thank the MIT BioMicro Center and the Babette Tang Histology Core, the Flow Cytometry Core, and the Microscopy Core at the Koch Institute of MIT technical support and instrumentation access. We greatly appreciate A. Shalek and the team at BIORAD, including D. Coe, A. Olcott, P. Schnepp, and A. Chowdhury, for an exciting collaboration on the snRNA sequencing portion of this project: BIORAD provisioned the snRNA sequencing reagents and provided technical advice for completion of these studies.

## Author contributions

AES, SNB, and CSC conceived this study. AES designed and analyzed all experiments. AES, VK, VH, JL, AJ, and AF contributed to experiments. CT and AES designed and performed the single nucleus RNA sequencing study. SNB and CSC supervised and funded the research. AES and CSC wrote this paper with input from all authors.

## Data and Materials Availability

All data needed to evaluate the conclusions in the paper are present in the paper and/or the Supplementary Materials. The single nucleus RNA sequencing data has been deposited at the Broad Single Cell Portal at under accession number SCP3418.

