## Supplemental Materials for "Synthetic Control of Implanted Engineered Liver Tissue Growth"

Amy E. Stoddard *et al.*

**This PDF file includes:**

Figs. S1 to S17

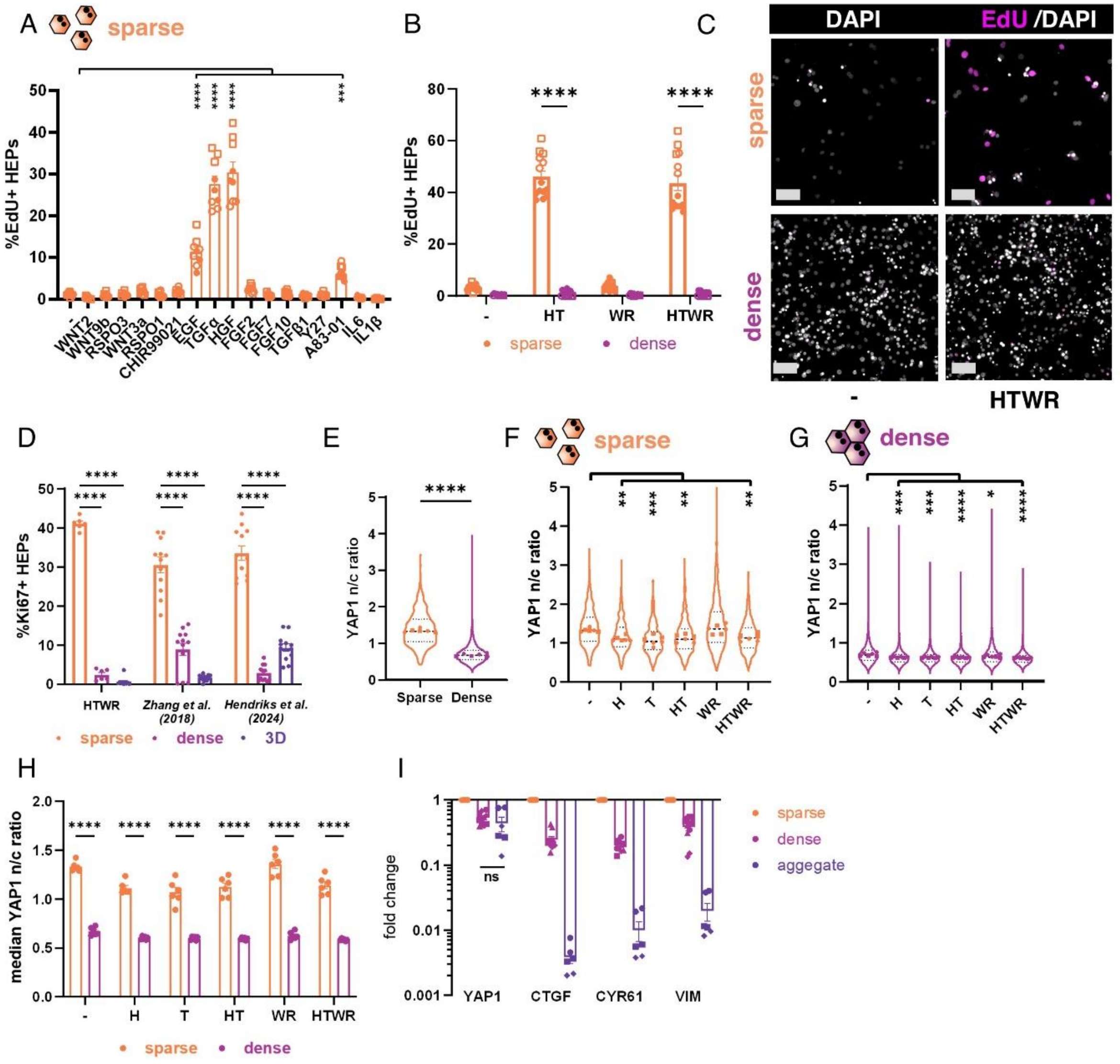


***Fig. S1. Density reduces YAP activity and proliferative response to GFs.*** **(A)** Proliferation of 2D sparse HEPs after 2 days of factor treatment in serum free media (N:3,n:3). **(B)** Proliferation of HEPs in response to recombinant GFs in sparse or dense 2D culture conditions (N:3,n:3). **(C)** Representative immunofluorescent images from B. Scale bars 100 µm. **(D)** Proliferation of HEPs treated with published HEP expansion medias in sparse, dense, or 3D cultures (N:3,n:3). **(E)** Quantification of YAP1 nuclear to cytoplasmic ratio in sparse/ dense cultures in basal media, **(F)** GF treated sparse cultures, and **(G)** GF treated dense cultures. Dots indicate median value of replicates. (N:2,n:3) **(H)** Median YAP nuclear to cytoplasmic ratio in 2D sparse and dense cultures, with and without GF supplementation (N:2,n:3). **(I)** YAP1 and YAP target gene RNA expression in HEPs cultures in sparse, dense, or aggregate cultures, fold change normalized to sparse cultures. All changes are significant p < .01 except for those marked (N:3,n:2). A,B,E,F,G,I: symbols denote data from separate experiments. Filled symbols in A,B denote experimental repeat shown in main figure. For ease of comparison, some data from Figure 1 is reproduced here. E,F,G: Statistics computed on medians of data set. (p *<.05, ** < .01, *** < .001, **** < .0001; A,F,G: one-way ANOVA, B,D,H,I: two-way ANOVA, E: two tailed t-test).

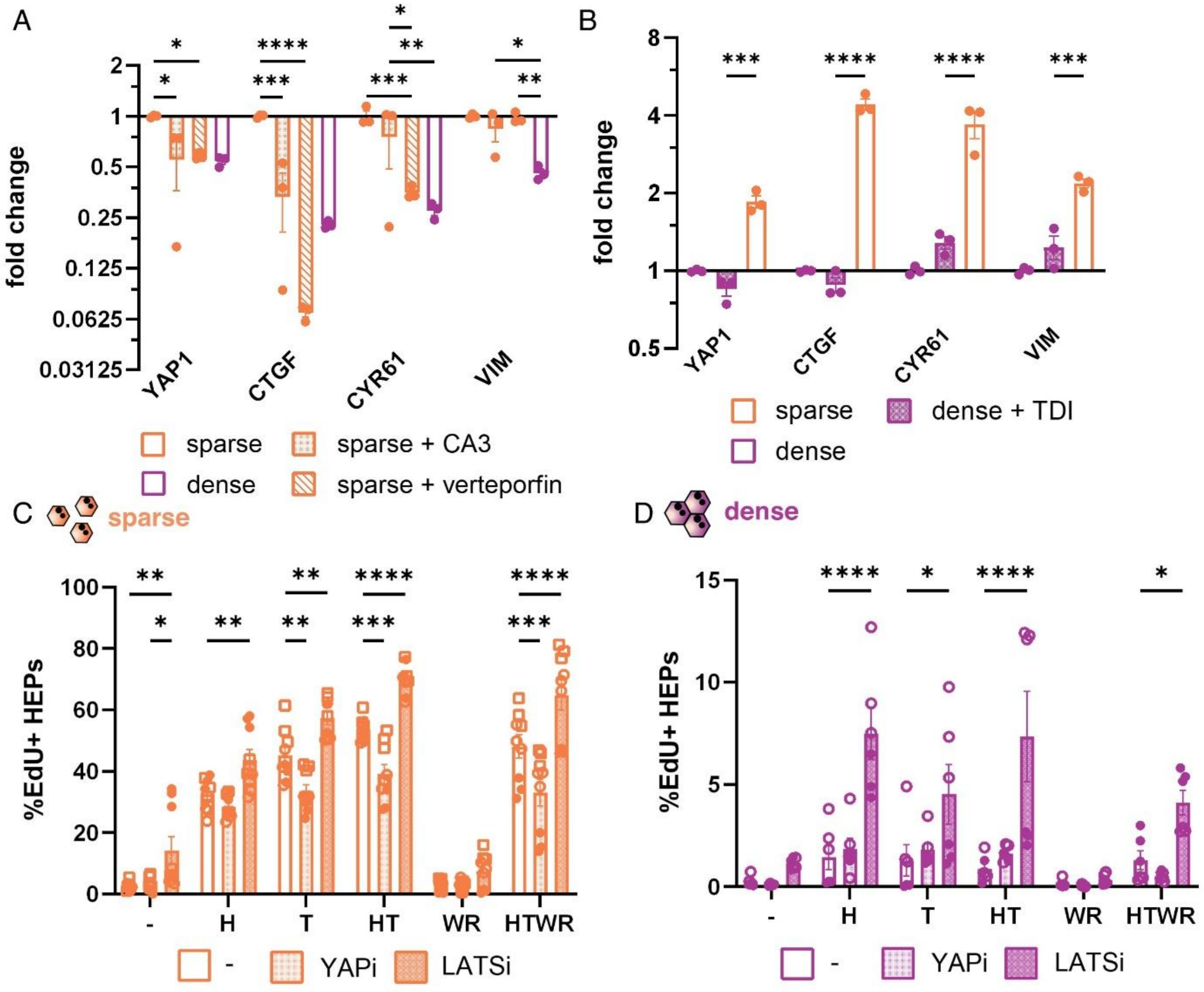


***Fig. S2. GF mediated proliferation is altered by YAP activity.*** **(A)** YAP1 and YAP target genes RNA expression in 2D HEP cultures with and without YAP inhibitors CA3 and verteporfin (N:1,n:3). **(B)** YAP1 and YAP target gene expression in 2D HEP cultures with and without LATS kinase inhibitor TDI-011536 **(N:1,n:3)**. **(C)** Proliferation of sparse and **(D)** dense cultures with and without treatment with growth factors, and YAP inhibitor CA3, or LATS inhibitor TDI-011536 (N:3,n:3). C,D: symbols denote data from separate experiments, filled symbols denote experimental repeat shown in main figure. (p *<.05, ** < .01, *** < .001, **** < .0001; A,B,C,D: two-way ANOVA).

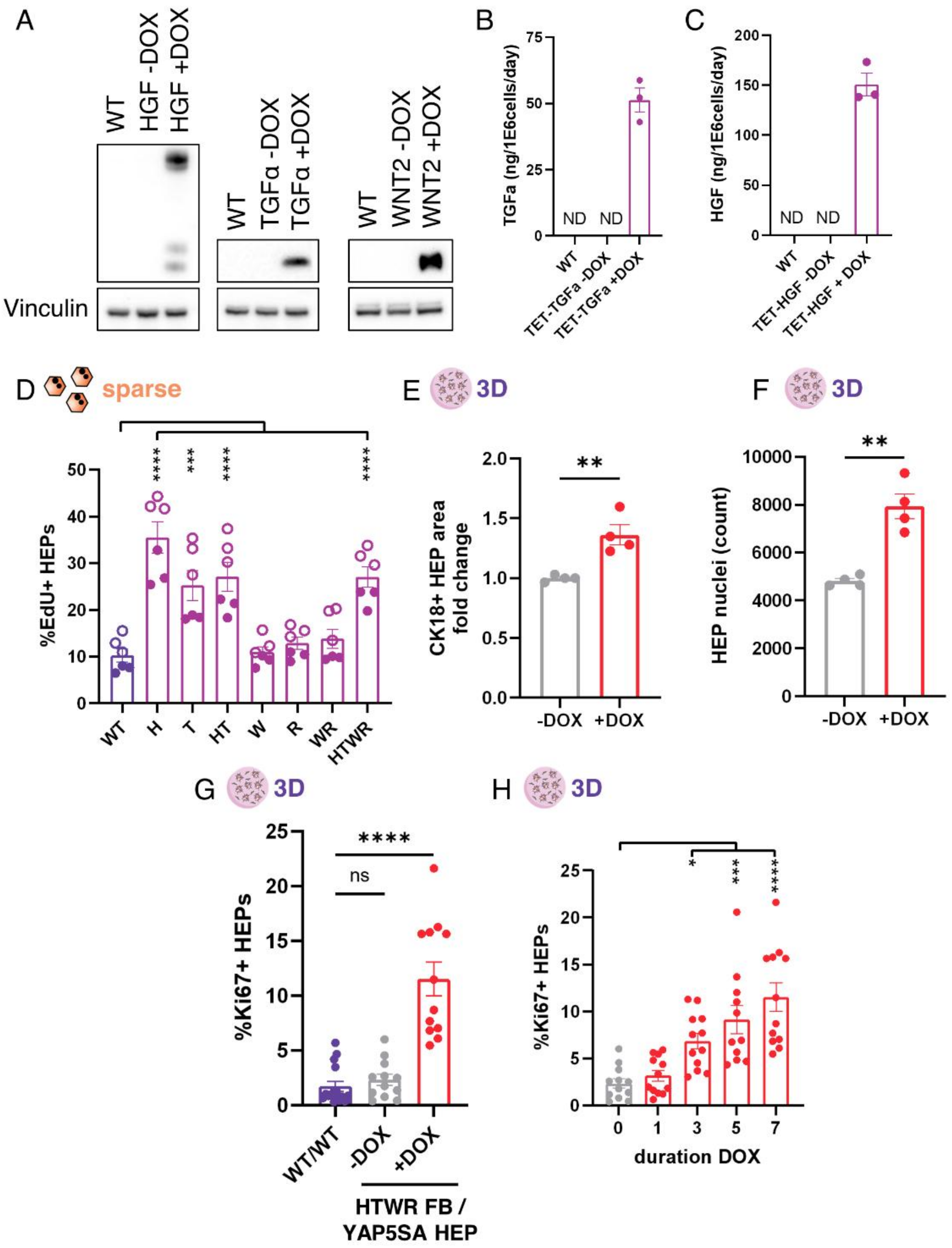


***Fig. S3. Synthetic control of GF and YAP signaling in 3D liver tissues.*** **(A)** Western blot of synGF FBs with and without DOX induction. **(B)** ELISA quantification of TGFα production from synGF FBs. **(C)** ELISA quantification of HGF production from synGF FBs. **(D)** Proliferation of 2D sparse HEPs treated with conditioned media from WT or synGF NHDFs for 2 days. Filled symbols denote experimental repeat shown in main figure (N:2,n:3). CK18+ HEP area **(E)** and HEP nuclei count **(F)** quantified from step sections of $FB^{HTWR}$ /$HEP^{YAP5SA}$ after 7 days of DOX induction compared to uninduced controls. **(G)** Proliferation of HEPS $FB^{HTWR}$ /$HEP^{YAP5SA}$ tissues induced with DOX compared to HEPs in uninduced and unengineered $FB^{WT}$/$HEP^{WT}$ tissues (N:3,n:4) **(H)** Proliferation of HEPs in $FB^{HTWR}$ /$HEP^{YAP5SA}$ tissues at day 7 after different duration DOX stimuli starting on day 0 (N:3,n:4). For ease of comparison, some data from Figure 2 is

***Fig. S3 cont.*** reproduced here. (p *<.05, ** < .01, *** < .001, **** < .0001; E,F: t-test, D,G,H: one-way ANOVA)

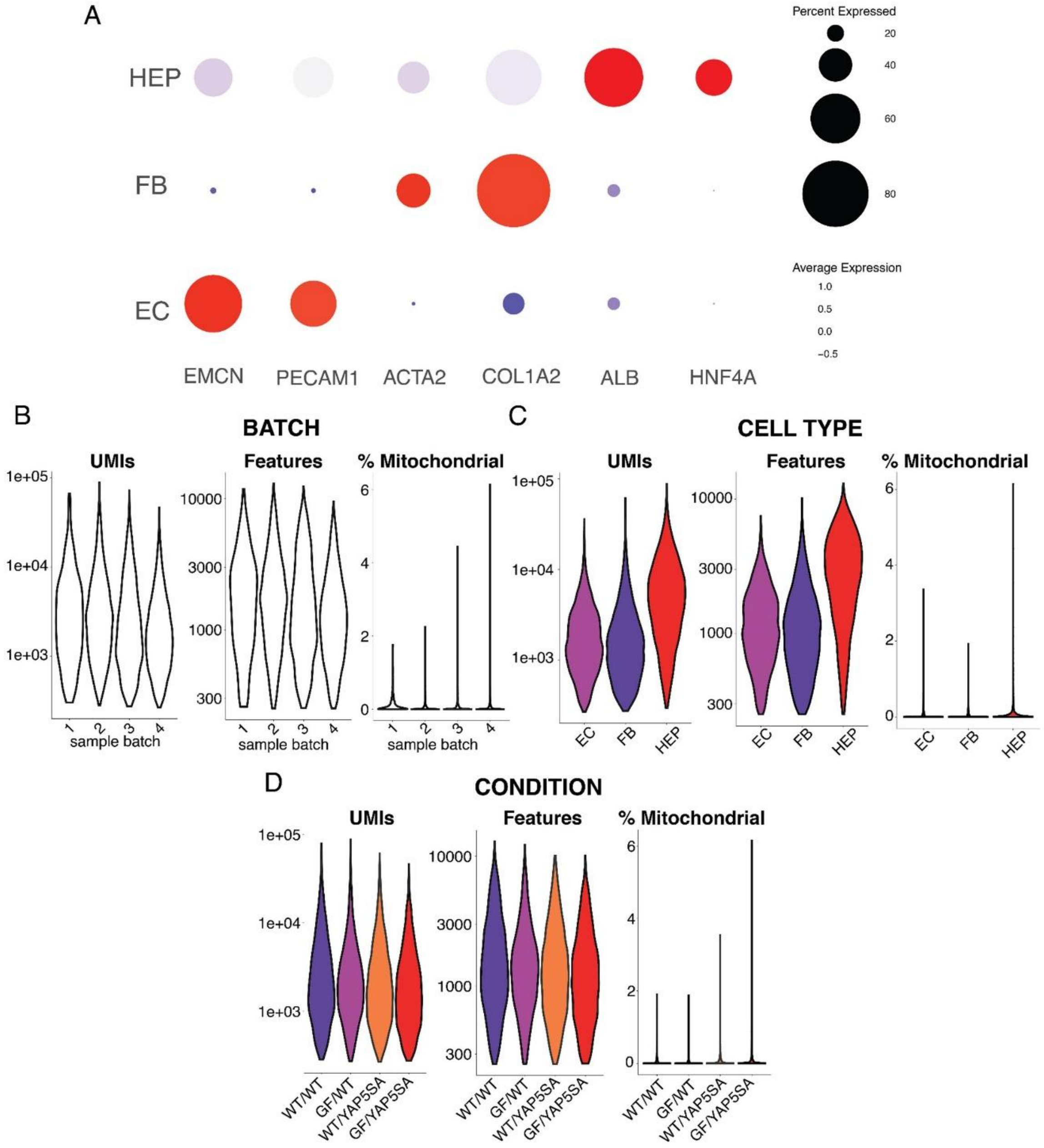


***Fig. S4. snRNA sequencing quality metrics.*** **(A)** Average expression of marker genes by EC, HEP, and FB clusters. QC metrics (RNA counts, feature counts, % mitochondrial RNA) by **(B)** batch, **(C)** cell type, and **(D)** experimental condition.

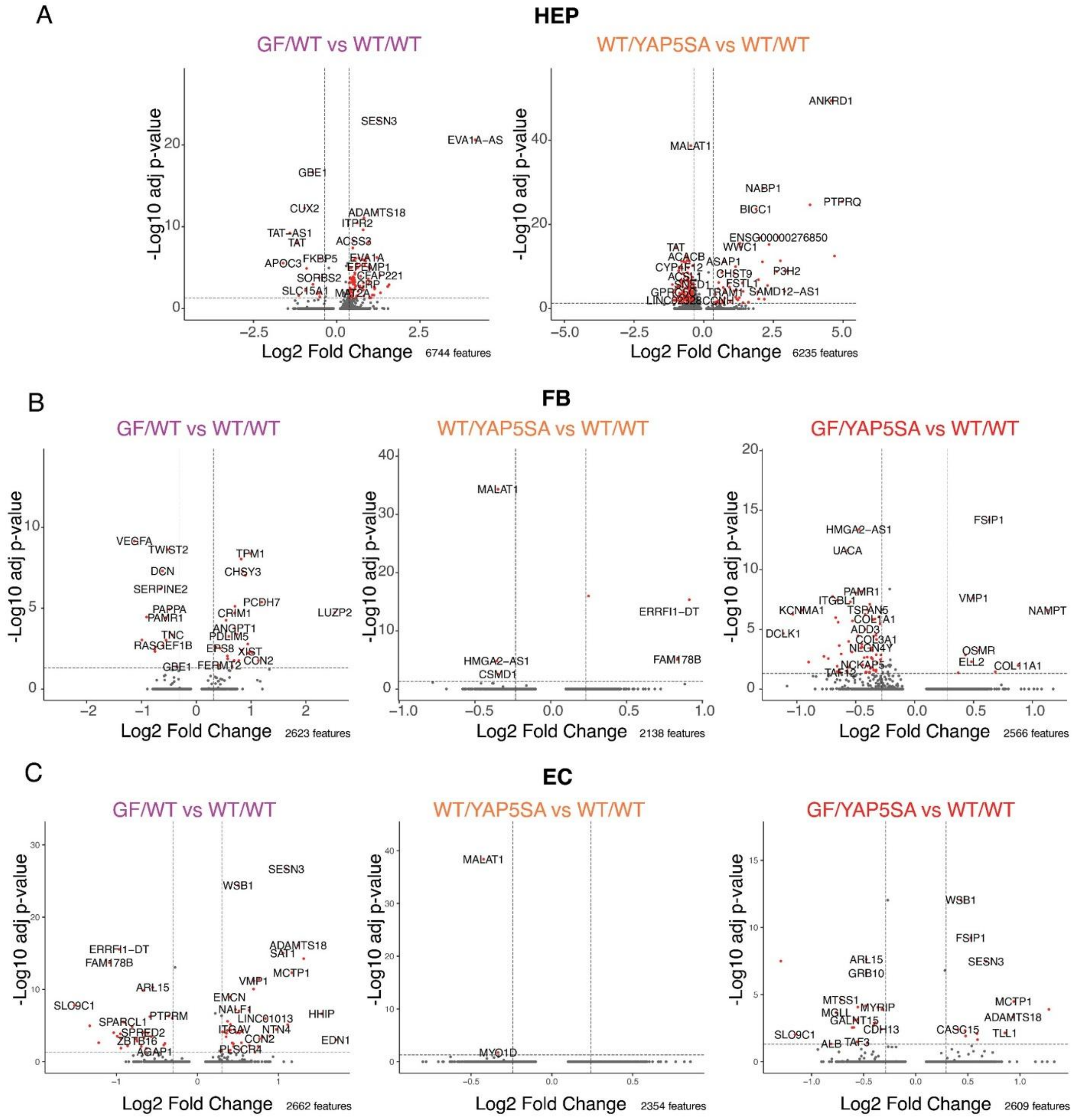


***Fig. S5. Differentially expressed genes in snRNA sequencing dataset.*** Volcano plots of differentially expressed genes in **(A)** HEP, **(B)** FB, and **(C)** EC cell populations compared to unengineered FB$^{WT}$/HEP$^{WT}$ controls.

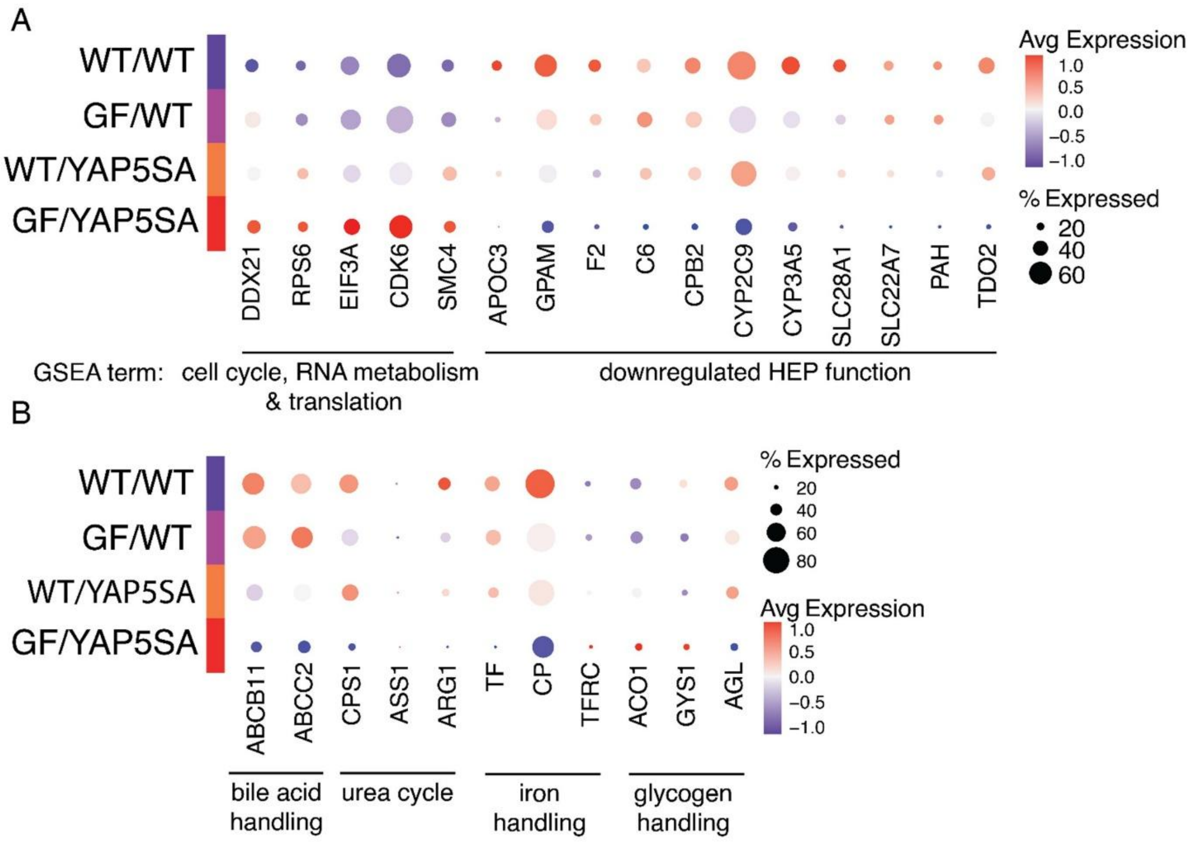


***Fig. S6. Differentially expressed genes associated with HEP proliferation and function.*** **(A)** Gene expression by condition of leading edge genes associated with perturbed gene sets shown in Figure 3C. **(B)** Gene expression by condition of genes associated with bile acid, urea cycle, iron, and glycogen functions of HEPs.

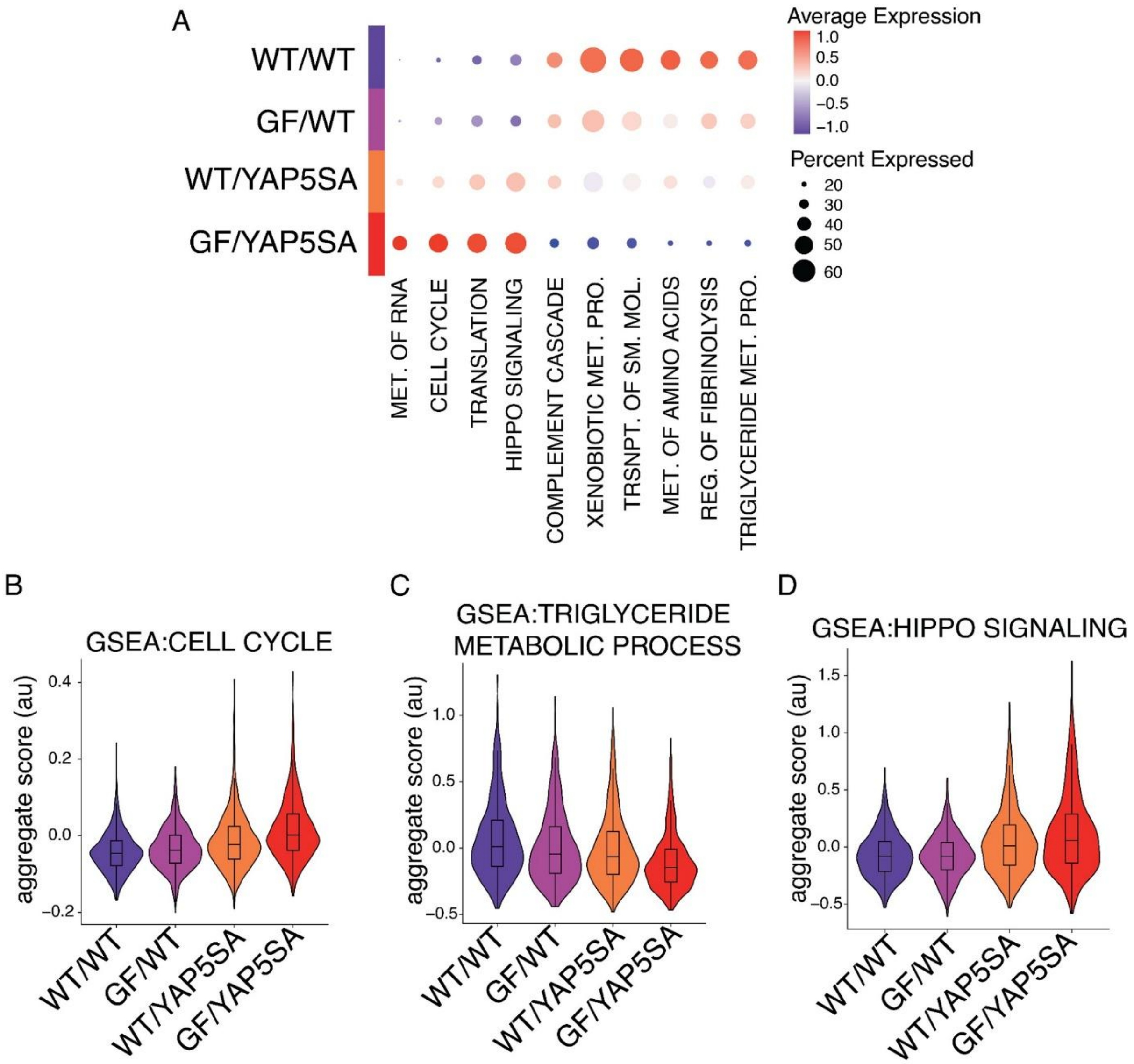


***Fig. S7. Dual synGF/YAP5SA activation most strongly changes HEP phenotype.*** **(A)** Module scoring across conditions on differentially expressed pathways shown in Figure 3C. Violin plots of **(B)** cell cycle, **(C)** triglyceride metabolism, and **(D)** hippo signaling genes set module scores by experimental condition.

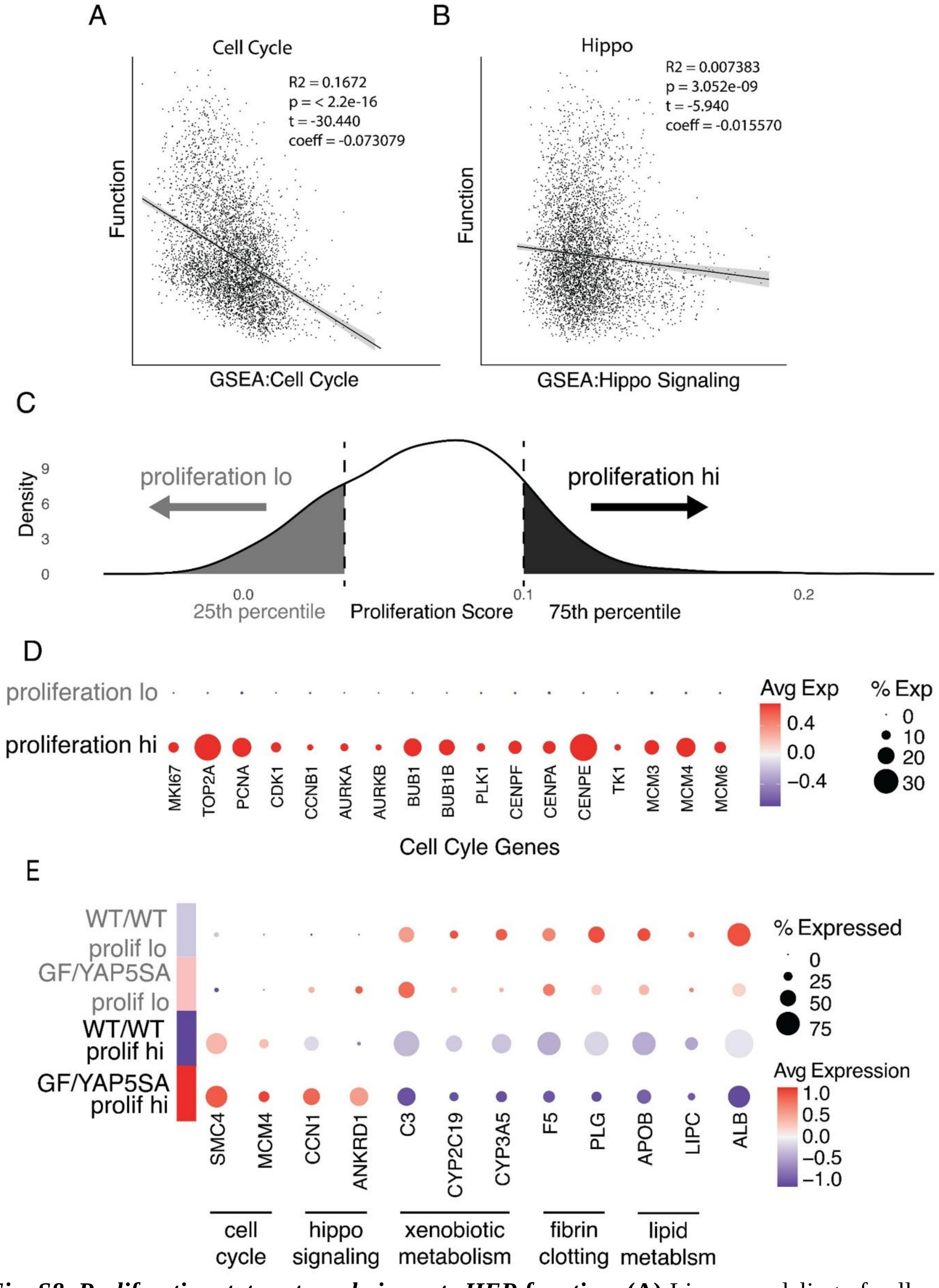


***Fig. S8. Proliferative status strongly impacts HEP function.*** **(A)** Linear modeling of cell cycle (proliferation) and **(B)** hippo signaling (YAP activity) vs function module scores of all HEPs, with

***Fig. S8 cont.*** goodness of fit statistics. **(C)** Graphical representation of HEP proliferation score binning, where “proliferation hi” HEPs are those in the top quartile of all HEPs across all conditions, and “proliferation lo” HEPs are those in the bottom quartile. **(D)** Expression of cell cycle associated genes in proliferation lo vs hi HEPs. **(E)** Expression of genes associated with pathways plotted in Fig 3I.

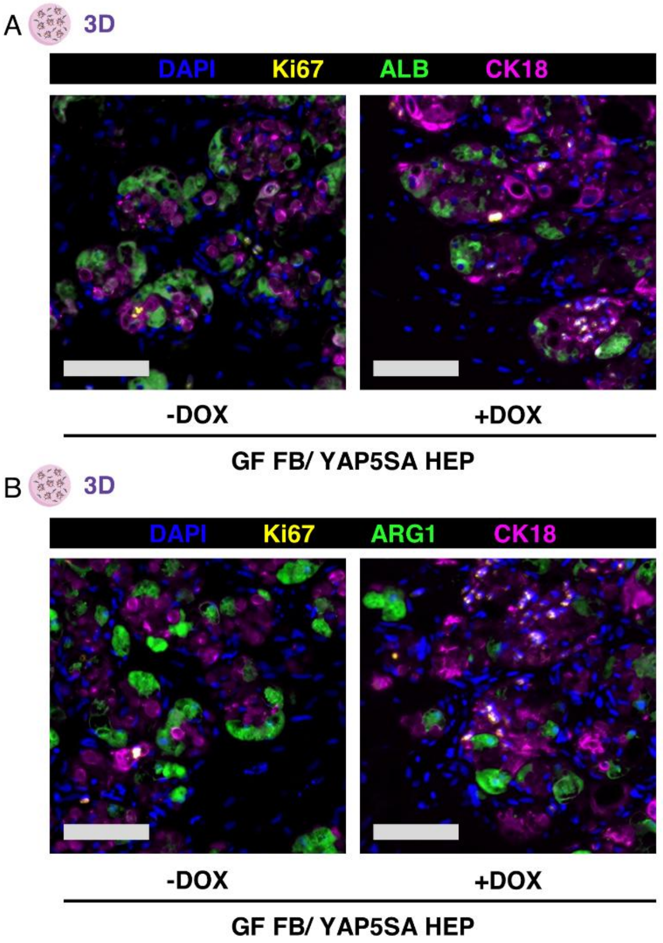


***Fig. S9. Proliferative HEPs express less functional proteins.*** **(A)** Immunohistochemical staining of ALB, Ki67 and HEP marker CK18 in FB$^{HTWR}$ /HEP$^{YAP5SA}$ *in vitro* liver tissues with or without 7 days of DOX induction. Scale bars 100 µm. **(B)** Immunohistochemical staining of ARG1, Ki67 and HEP marker CK18 in FB$^{HTWR}$ /HEP$^{YAP5SA}$ *in vitro* liver tissues with or without 7 days of DOX induction. Scale bars 100 µm.

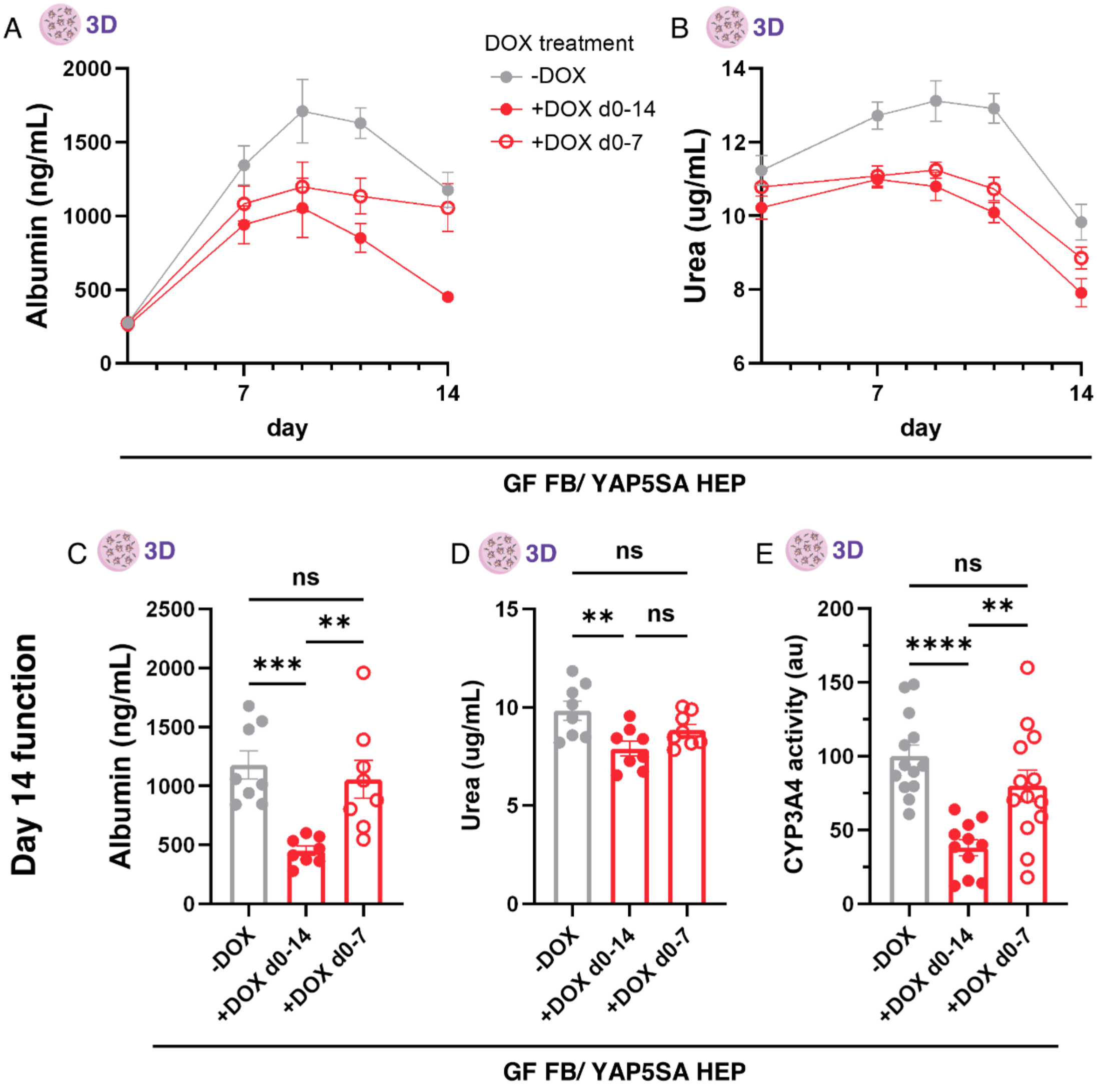


***Fig. S10. Kinetics of engineered liver tissue function.*** Kinetic time course of **(A)** albumin and **(B)** urea production by $FB^{HTWR}$ /$HEP^{YAP5SA}$ tissues uninduced or induced with DOX from day 0-7 or from day 0-14. Day 14 quantification of (C) albumin, (D) urea, and (E) CYP3A4 activity measure in $FB^{HTWR}$ /$HEP^{YAP5SA}$ tissues uninduced or induced with DOX from day 0-7 or from day 0-14. (p ** < .01, *** < .001, **** < .0001; C,D,E: one-way ANOVA)

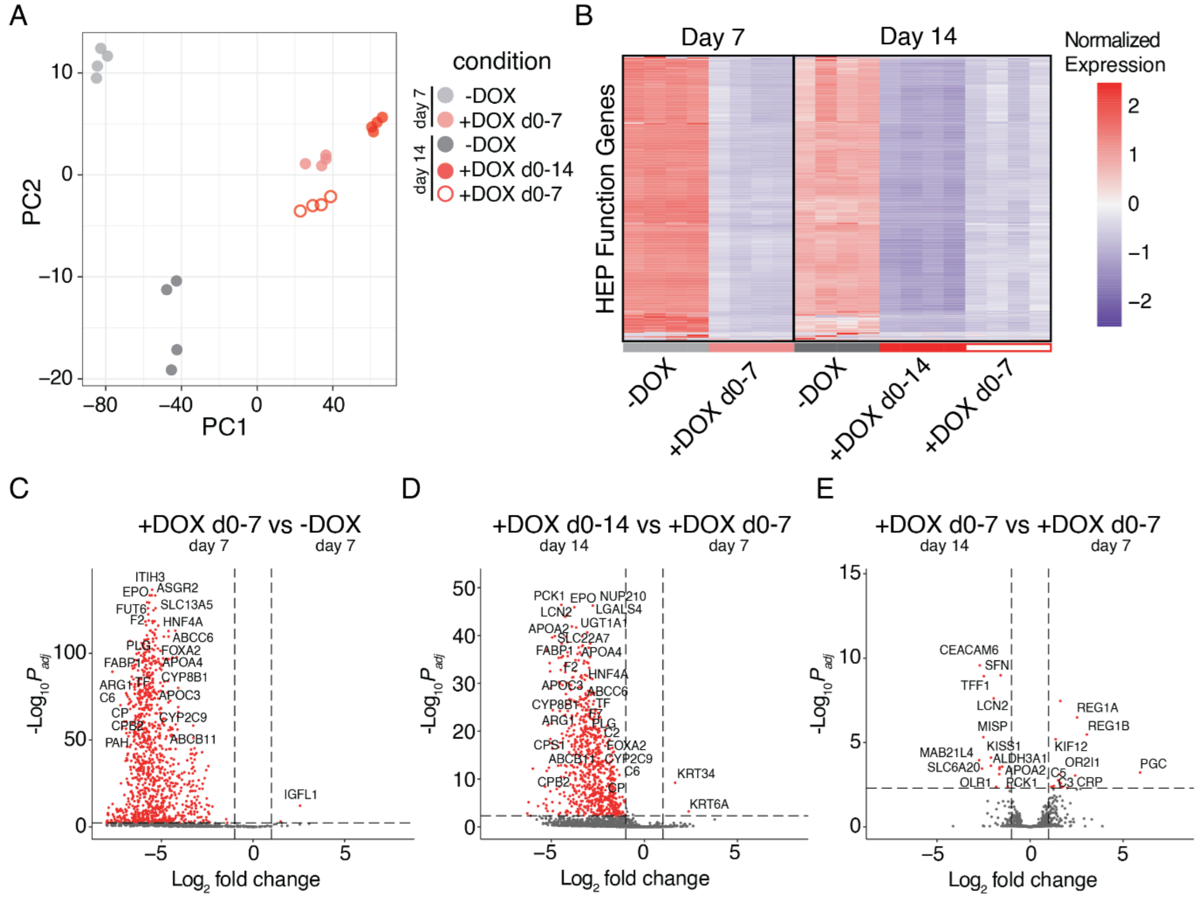


***Fig. S11. Bulk sequencing analysis of HEP function in FB$^{HTWR}$ /HEP$^{YAP5SA}$ tissues by DOX duration.*** **(A)** PC clustering of samples based on normalized expression of HEP specific genes. PC1 ~ 98% variance, PC2 ~ 1.5% variance. **(B)** Heatmap of normalized expression of 695 HEP function related genes by day of isolation (day 7, day 14) and duration of DOX treatment. **(C)** Differentially expressed genes upon induction of DOX for 7 days. **(D)** Genes differentially expressed when tissues treated for an additional 7 days of DOX. **(E)** Genes differentially expressed after 7 days of DOX withdrawal compared to pre-withdrawal 7 day DOX induction baseline.

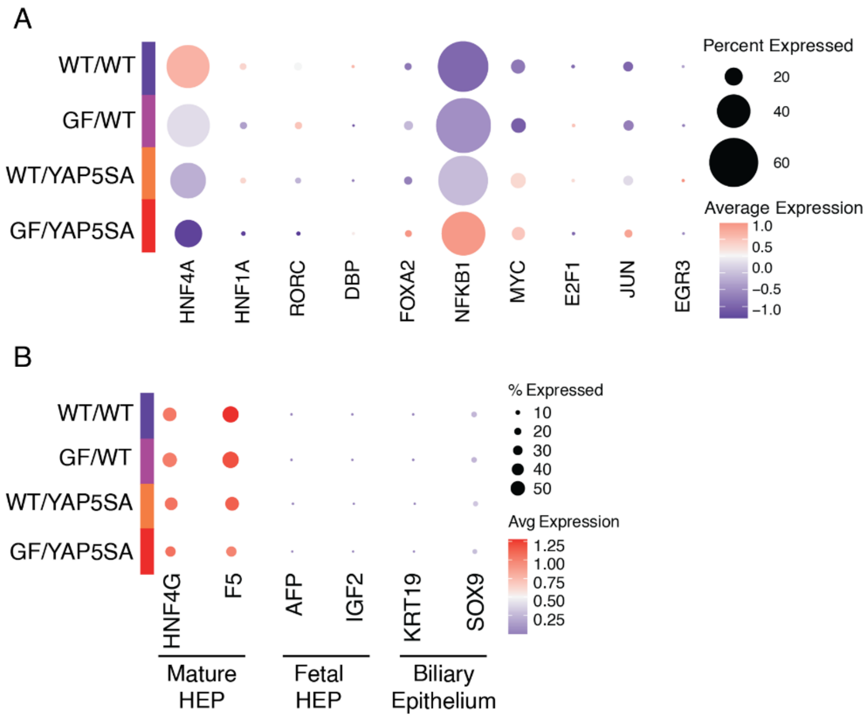


***Fig. S12. Transcription factor, fetal and biliary markers.*** **(A)** Expression of transcription factors with predicted altered activity. **(B)** Marker gene expression for mature HEPs, fetal HEPs, and biliary epithelial cells.

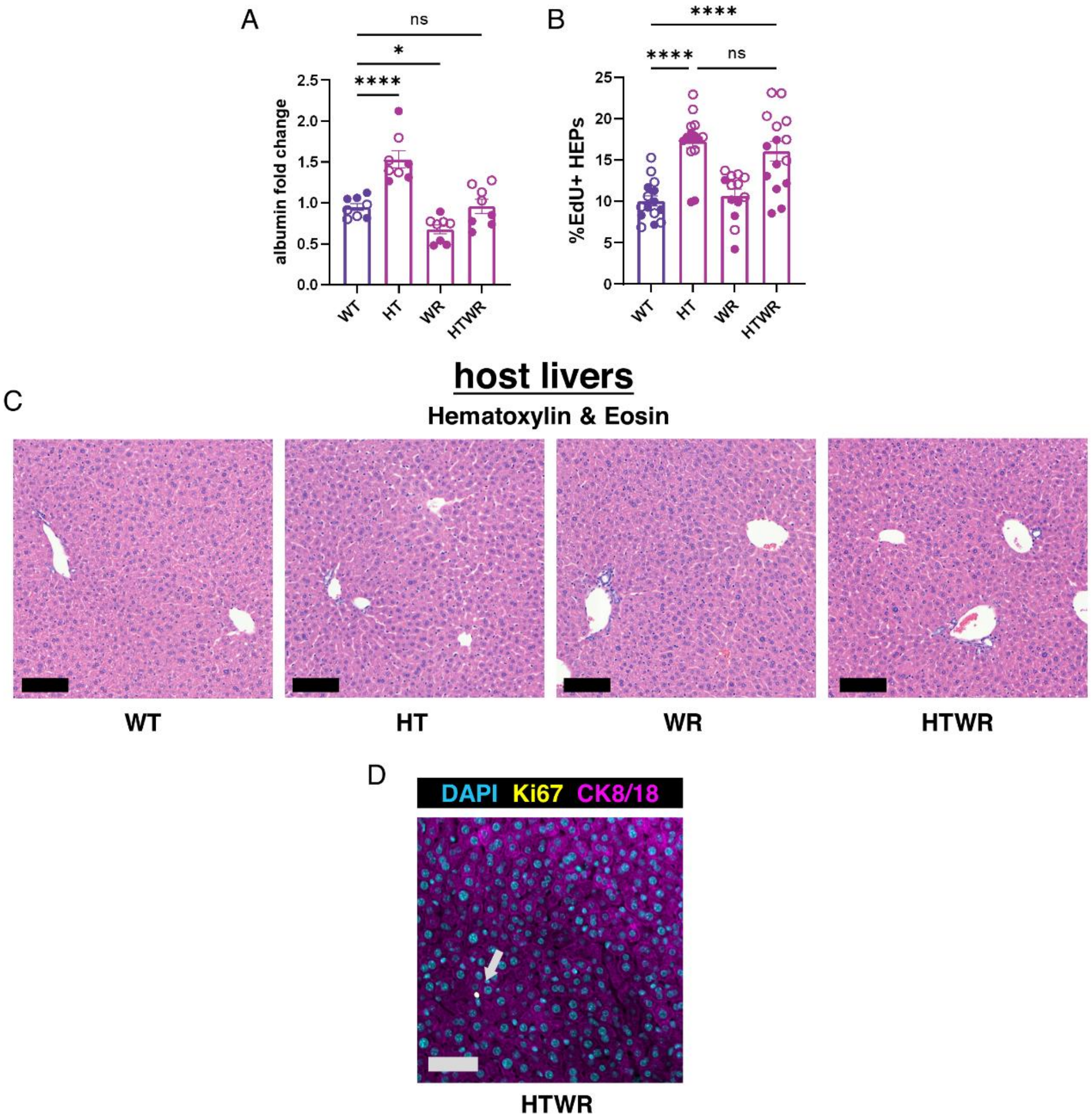


***Fig. S13. Synthetic expression of GFs in implanted liver tissues.*** **(A)** Serum human albumin fold change compared to preinduced baseline in mice implanted with ectopic liver grafts after 1 week of synGF expression (N:8,n:2). **(B)** Quantification of HEP proliferation during 1 week in vivo induction of synGFs in implanted liver tissues (N:8,n:2). Filled symbols denote experimental repeat shown in Figure 4B and is reproduced here for ease of comparison. **(C)** H&E staining of mouse host livers after 1 week of expression of synGFs from implanted ectopic liver tissues. Scale bars 100 µm. **(D)** HEPs in mouse host livers from animals implanted with synGF expressing ectopic liver tissues were not more proliferative than controls. Only rare Ki67+ nonparenchymal cells were observed (grey arrow). Scale bars 50 µm.

A

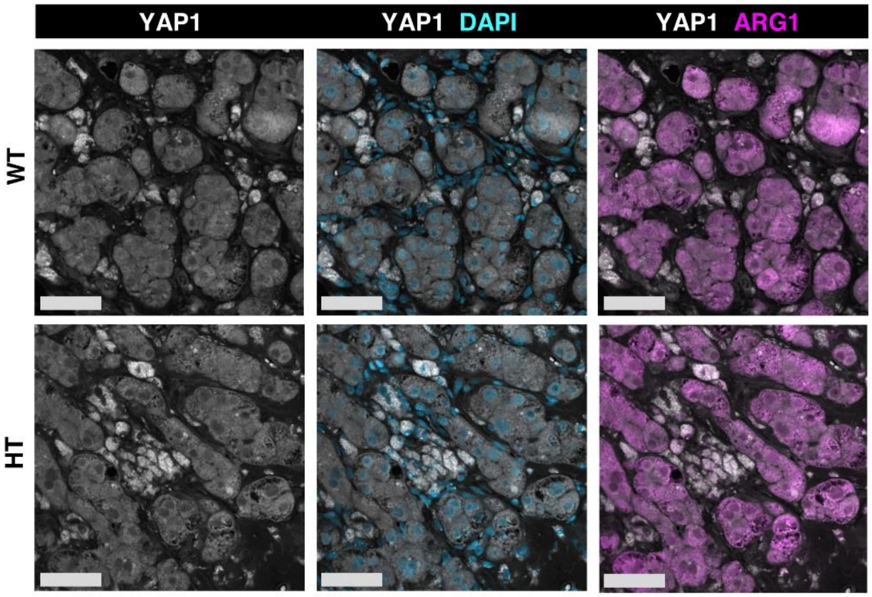


***Fig. S14. YAP is excluded from the nucleus in implanted ectopic liver tissues.***
**(A)** Immunohistochemical staining of YAP1 and HEP marker arginase 1 in WT or synHT expressing ectopic liver tissues. Some images are reproduced from main figure for ease of comparison. Scale bars 50 µm.

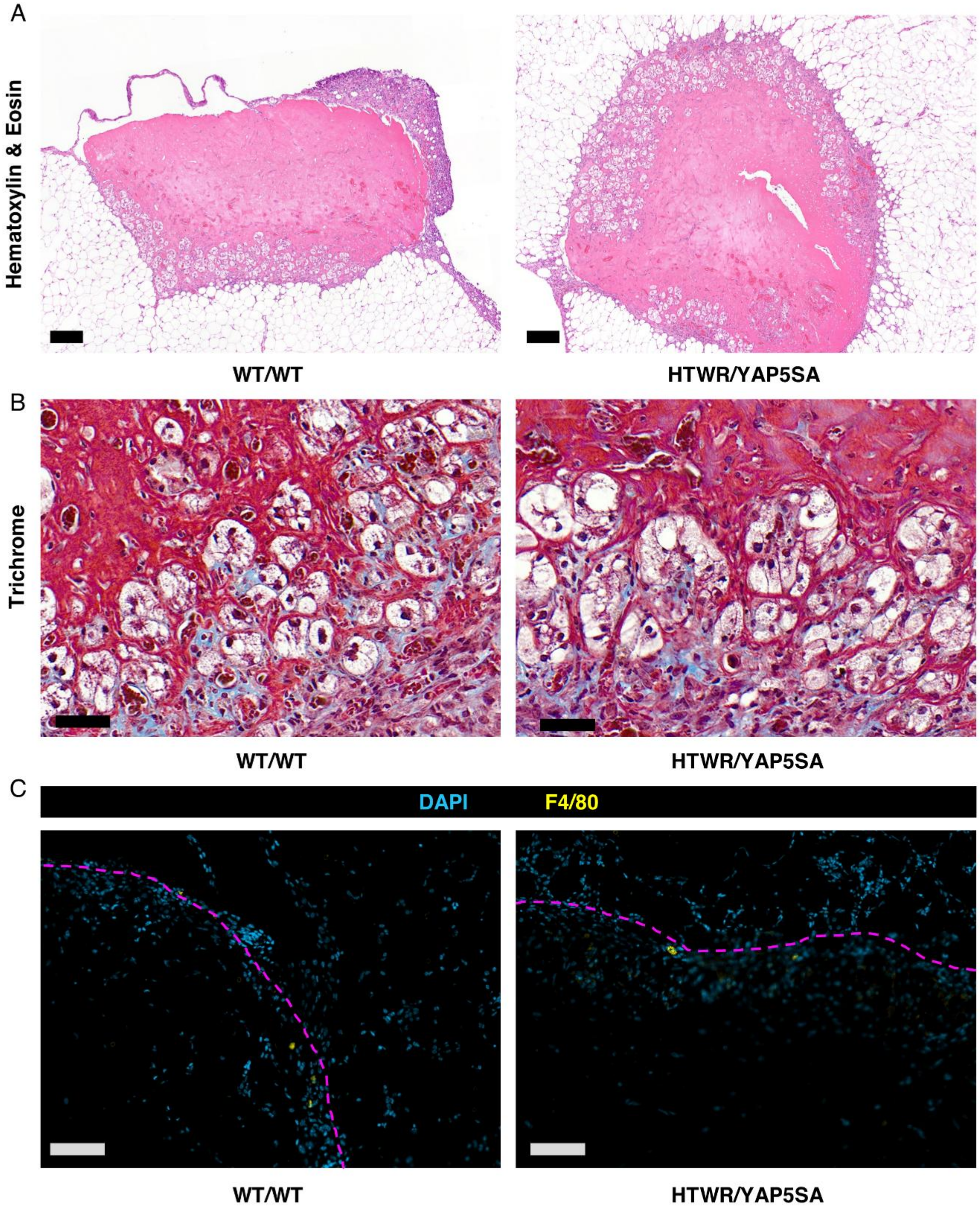


***Fig. S15. Histological assessment of implanted FB$^{HTWR}$/HEP$^{YAP5SA}$ tissues after 1 week of DOX induction.*** **(A)** H&E staining of ectopic liver tissues at 1 week post implantation. Scale bars 100 µm. **(B)** Modified Masson's Trichrome staining of ectopic liver tissues at 1 week post implantation, where collagen is blue and nuclei are black. Scale bars 50 µm. **(C)** Staining of macrophage marker F4/80. Graft boundaries denoted by magenta dotted line. Scale bars 200 µm.

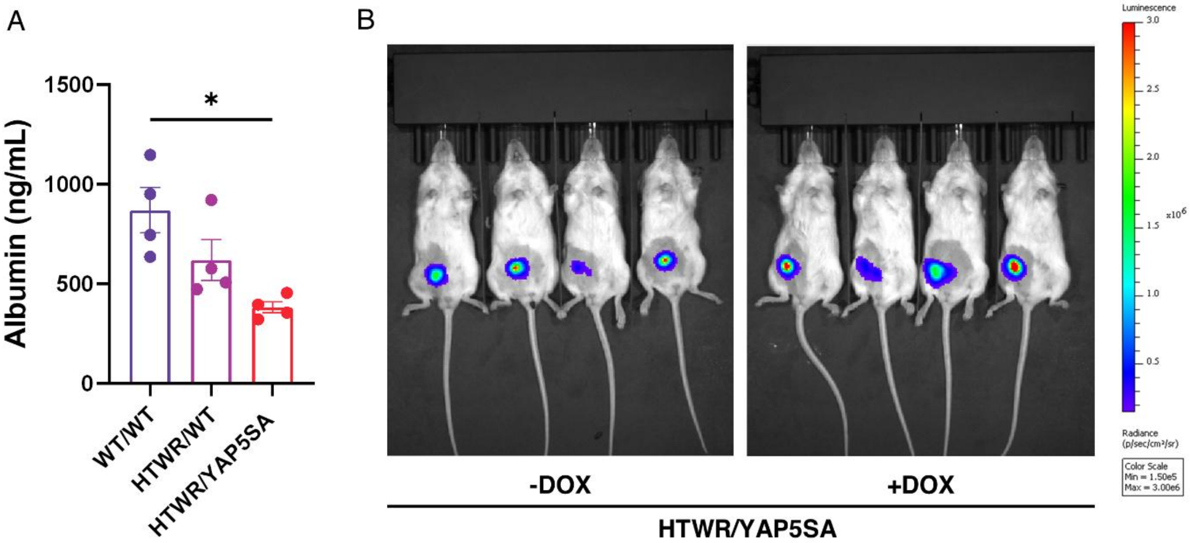


***Fig. S16. Function and expansion of implanted FB$^{HTWR}$ /HEP$^{YAP5SA}$ tissues after 1 week of DOX induction.*** **(A)** Serum human albumin from animals implanted with FBWT /HEPWT, FBHTWR /HEPWT, or FBHTWR /HEPYAP5SA ectopic liver tissues measured after 1 week of DOX induction (N:6,n:1). **(B)** Luminescent signal from reporter YAP5SA expressing HEPs quantified in Figure 4I, with and without DOX induction.

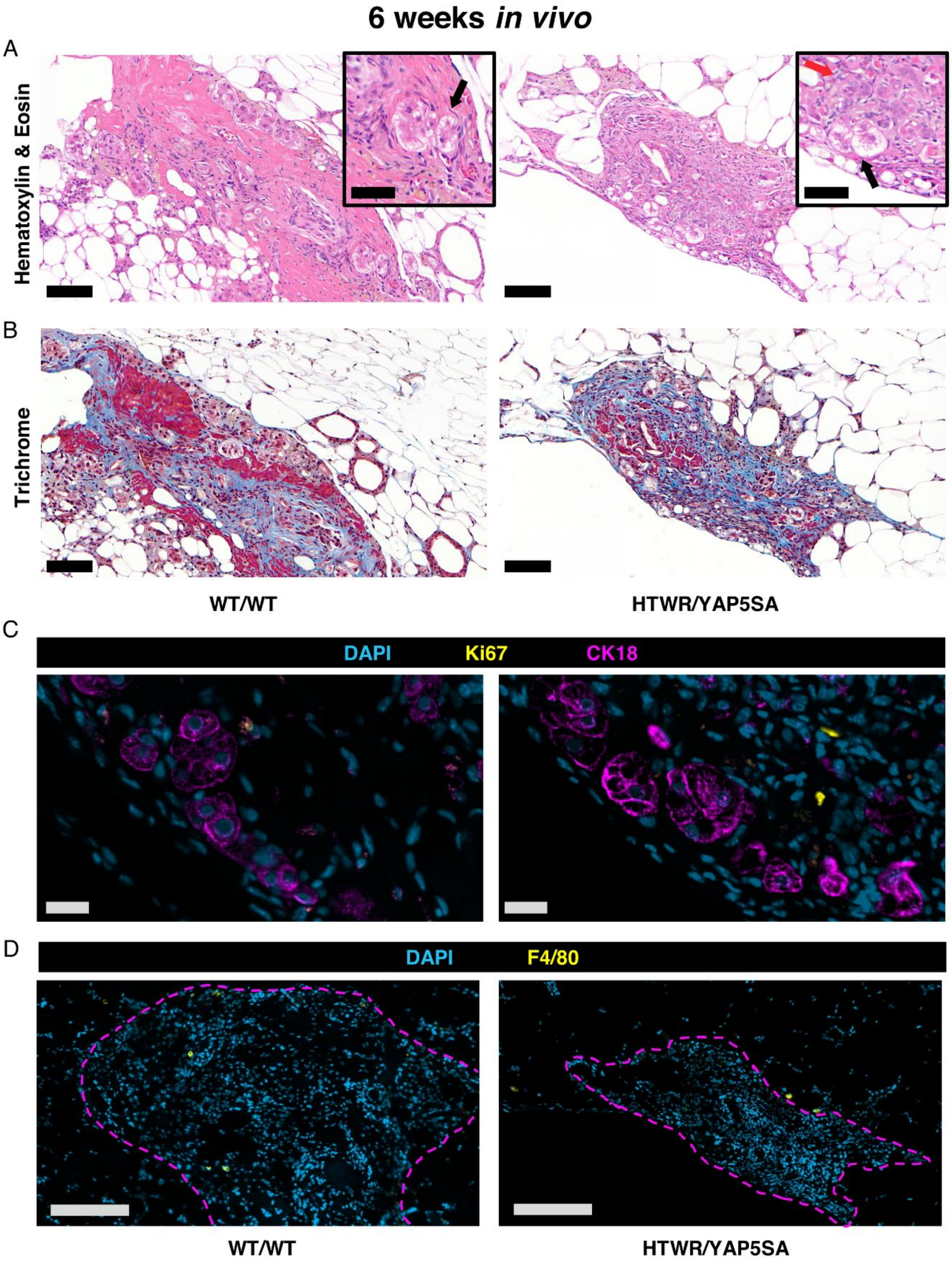


***Fig. S17. Histological assessment of implanted FB$^{HTWR}$/HEP$^{YAP5SA}$ tissues after 6 weeks of DOX induction.*** **(A)** H&E staining of ectopic liver tissues at 6 weeks post implantation. Scale bars 100 µm. **(B)** Modified Masson's Trichrome staining of ectopic liver tissues at 6 weeks post

***Fig. S17 cont.*** implantation, where collagen is blue and nuclei are black. Scale bars 50 µm. **(C)** Staining for Ki67 and HEP marker CK18. Scale bars 20 µm. **(D)** Staining of macrophage marker F4/80. Graft boundaries denoted by magenta dotted line. Scale bars 200 µm.
